# A Wi-Fi live streaming Centrifuge Force Microscope for benchtop single-molecule experiments

**DOI:** 10.1101/2020.07.07.192393

**Authors:** Jibin Abraham Punnoose, Andrew Hayden, Lifeng Zhou, Ken Halvorsen

## Abstract

The ability to apply controlled forces to individual molecules has been revolutionary in shaping our understanding of biophysics in areas as diverse as dynamic bond strength, biological motor operation, and DNA replication. However, the methodology to perform single-molecule experiments has been and remains relatively inaccessible due to cost and complexity. In 2010, we introduced the Centrifuge Force Microscope (CFM) as a new platform for accessible and high-throughput single-molecule experimentation. The CFM consists of a rotating microscope where prescribed centrifugal forces can be applied to microsphere-tethered biomolecules. In this work, we develop and demonstrate a next-generation Wi-Fi CFM that offers unprecedented ease of use and flexibility in design. The modular CFM unit fits within a standard benchtop centrifuge and connects by Wi-Fi to a external computer for live control and streaming at near gigabit speeds. The use of commercial wireless hardware allows for flexibility in programming and provides a streamlined upgrade path as Wi-Fi technology improves. To facilitate ease of use, detailed build and setup instructions are provided, as well as LabVIEW™ based control software and MATLAB^®^ based analysis software. We demonstrate the analysis of force-dependent dissociation of short DNA duplexes of 7, 8, and 9 bp using the instrument. We showcase the sensitivity of the approach by resolving distinct dissociation kinetic rates for a 7 bp duplex where one G-C base pair is mutated to an A-T base pair.

**Significance:** The ability to apply mechanical forces to individual molecules has provided unprecedented insight into many areas of biology. Centrifugal force provides a way to increase the throughput and to decrease the cost and complexity of single-molecule experiments compared to other approaches. In this work, we develop and demonstrate a new user-friendly Centrifuge Force Microscope (CFM) that enables live-streaming of high-throughput single-molecule experiments in a benchtop centrifuge. We achieved near gigabit bandwidth with standard Wi-Fi components, and we provide detailed design instructions and software to facilitate use by other labs. We demonstrate the instrument for sensitive kinetic measurements that are capable of resolving the difference between two DNA duplexes that differ by a single G-C to A-T substitution.

## Introduction

Force-based single-molecule techniques are powerful approaches to understand complex reaction pathways and dynamic interplay of biomolecules. Techniques such as optical and magnetic tweezers and atomic force microscopy (AFM) are the most widely used force-based single-molecule techniques, and have contributed to our understanding of the influence of force on biological structures and processes (1–4). Despite the success of these methods, they are typically complex, costly, and have low throughput due to serial probing of one molecule at a time.

The Centrifuge Force Microscope (CFM) was introduced to address these problems and was the first method to demonstrate thousands of parallel single-molecule experiments (5). Conceptually, the CFM is a rotating microscope that applies centrifugal forces to many singlemolecule tethered particles while tracking their microscopic motions with a camera (Figure 1a & 1b). Since its introduction, the CFM has evolved substantially. The first generation of CFM was a relatively crude, custom built open-air centrifuge with an attached microscope unit (Figure 1c). While it achieved the goal of demonstrating massive multiplexing in single-molecule pulling experiments (achieving >5000 simultaneous experiments), the prototype had many shortcomings including safety and difficulty in design, assembly and operation. Several of these issues were solved in the second generation, which used a custom modified commercial centrifuge to contain the microscopy module (6, 7). The third generation enabled use of an off-the-shelf benchtop centrifuge by making the CFM a completely wireless plug-and-play module that fit in a standard centrifuge bucket (8). Outside of the 2 labs involved in the initial CFM development, a few labs have developed instruments on their own with various features. The Forde lab developed a mini-radio CFM that used wireless radio frequency transmission for a plug-and-play CFM module (9). The Hu lab developed a CFM that incorporated digital holography for 3D particle tracking (10).

**Figure 1:**
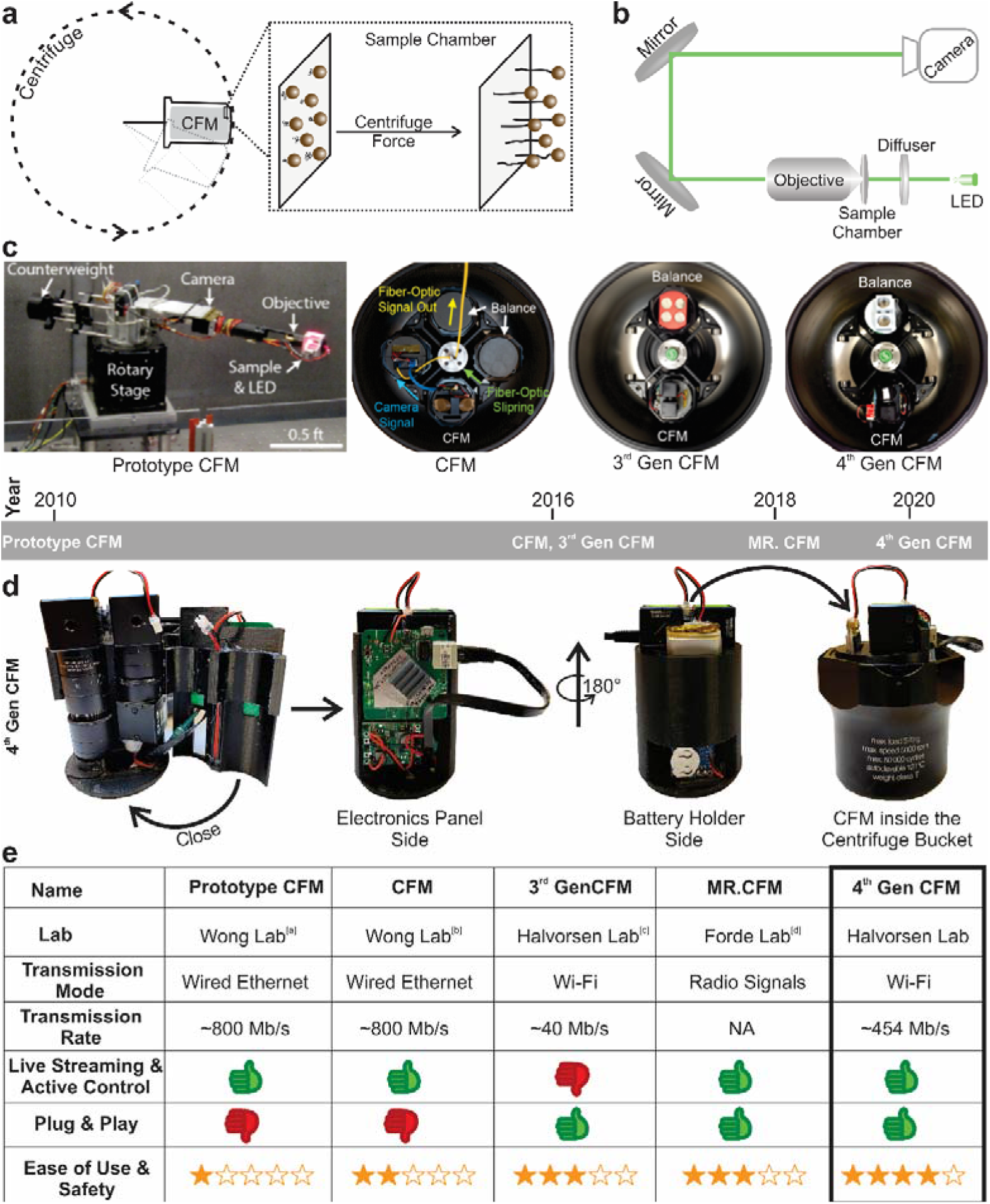
Working principles of Centrifuge Force Microscope and its evolution. (a) Schematics of working principle of CFM. Force is applied to the molecule of interest tethered between a glass-slide and microscopic bead, placed inside a centrifuge. The force experienced by the sandwiched molecule is propotional to the square of the angular velocity and the size of the bead. (b) Schematics of the optic assembly of CFM. The tetherd microsphere is visualized using a 40X or 20X infinite conjugate objective and imgaged using a machine vision camera. (c) Pictures of CFM belonging to various generation starting from the prototype CFM to the 4^th^ generation of CFM (current work). Pictures of CFMs developed outside Wong/Halvorsen lab is not shown. (d) Picture of 4^th^ generation CFM from various angles. (e) Table of properties comparing different versions of CFMs. Current version of CFM reported in this paper retains best featues from all the generations.

While the many implementations of the CFM have been generally evolving in the right direction, the designs so far still face tradeoffs between complexity and performance. In a key shortcoming, our third generation CFM was wireless and plug-and-play, but lacked live streaming and real-time control. Here, we report development of our fourth generation CFM (Figure 1d), a plug-and–play device with real-time device control and data transmission over Wi-Fi. This device can be used in a commercial centrifuge without any customization and includes a user friendly interface that could lead to increased accessibility of this technique by other labs. Using this next generation CFM, we demonstrate single-molecule DNA shearing experiments investigating the effect of force, length and GC content on dissociation kinetics.

## Materials and Methods

### Instrument design and construction

The CFM module, designed to fit within a 400 mL bucket of a commercial centrifuge (Sorvall X1R), has aspects of similarity with our previous design (8). As with our previous designs, we put emphasis on reproducibility, and provide detailed parts lists, build instructions, and 3D models in the supplementary materials. The optical CFM components are largely unchanged from previous versions: a 40X infinity corrected objective, turning mirrors, LED and a diffuser. The main change in optical components was the camera, which was changed from a USB camera to a gigabit Ethernet camera (www.flir.com, Model # BFLY-PGE-50H5M-C) to facilitate the Wi-Fi capabilities. The optical arrangement and housing have only minor changes detailed in the Figure S1.

The electrical system and the mechanical housing to mate with the centrifuge bucket were redesigned for the Wi-Fi strategy, with slightly different designs for each of the two routers we used. For power distribution, a replaceable Lithium ion battery (www.adafruit.com, part # 2011) is connected to independent 5V and 12 V voltage adapters (www.pololu.com, part # 2891, 2895) to power the Wi-Fi router (www.tp-link.com, TL-WR902AC) and camera, respectively. Wiring diagrams and sketches of the instruments using two different routers are provided (Figures S2-S5).

We assembled the required electronics including the voltage adaptors for the camera and router, and an LED light source on a 3D printed housing (Figure 1d, Figure S3, S5, 3D files for housing are provided as supplementary files 1-6). All the parts required for building the CFM is listed in Table S1.

We also have designed a counterbalance, mass of which can be adjusted by adding or removing quarters. To maintain the center of mass, the counter balance was designed to resemble the shape of the fully assembled CFM, printed with 40 percent infill with hollow cylinder bore in the place of lens tube where quarters were added to fine tune the mass. A detailed protocol for making counter balance is described in the supplementary note 1 (Figure S6).

### Software and Networking

The housing for the CFM components and counter balance were designed using Fusion360 2019 (www.autodesk.com). The 3D files generated for the components are given as supporting files. The 3D files were converted to printer-specific g-code using the cura v3.4.1 (ultimaker.com).

The CFM controller was written using LabVIEW™ 2018 (www.ni.com). NI-IMAQdx driver in the vision acquisition module can communicate with GigE camera and programs can be written with built-in functions. The programs written for controlling FLIR Blackfly (Cat# BFLY-PGE-50H5M-C) with firmware version v1.42.3.00 is given as a supplementary file 7, with the front panel shown in Figure S7. To achieve wireless control of the camera using GigE drivers in LabVIEW™, a virtual ethernet connection was established by bridging the Wi-Fi adaptor and the ethernet adaptor. A detailed procedure is given in the supplementary note 2 and 3. To achieve good signal reception, we used a desktop PCI-e network adaptor (Intel 6050) with external antennas. To ensure the best data transmission, we activated the “resend lost packets” feature on the camera (specific parameters reported in Table S2).

### Sample and microsphere preparation

DNA constructs were prepared by hybridizing 123 oligonucleotides (Integrated DNA Technologies) to 7249 nt single-stranded M13mp18 DNA, as described in the previous work (8). The oligo hybridizing to the 3’ end of the M13 DNA contains double biotin on its 5’ end for immobilization to streptavin coated glass surface or the bead. The oligo hybridizing to the 5’ end of the M13 DNA extends beyond the M13, resulting in a 3’ single-stranded overhang (Figure S8). Different constructs can be prepared by just varying this oligo hybridizing the 5’ end of M13 DNA. For shearing experimets in this paper, the construct pairs that are immobilized to the glass slide and to the microspheres contains the 5’ hybridizing oligonucleotide that has complementary sequence in the overhang region. The list of all oligos used is given in Table S3.

To immobilize above prepared DNA constructs to streptavidin coated microspheres (Thermo Fisher Dynabeads M-270 2.8 μm diameter), we took 20 μl of streptavidin microspheres and washed thrice with 50 μl of phosphate-buffered saline (PBS) + 0.1% Tween 20. Following the wash, the beads were brought to a 10 μl volume, and 10 μl of the DNA construct (~500 pM) was added to it and shaken in a vortexer for 30 minutes. The unbound DNA was removed by washing the beads thrice with 50 μl of PBS + 0.1% Tween 20. The beads were brought to 40 μl volume.

### Chamber preparation

The reaction chamber is prepared according to previous work (8). Briefly, the reaction chamber consist of an 18mm and a12mm circular microscope glass slide (Electron Microscopy Sciences, catalogue # 72230-01 & 72222-01) sandwiching two parallel strips of Kapton tape (www.kaptontape.com) creating a channel of ~ 2 mm between the glass-slides. The glass chamber is assembled on top of a SM1A6 theread adaptor (Thor Labs). Streptavidin (Amresco) was passively adsorbed to the surface by passing 10 μl of streptavidin (0.1 mg/ml) in 1X PBS. After 1 minute of incubation, the chamber was washed thrice with 50 μl of PBS + 0.1% Tween 20 to remove any unbound streptavidin. Next, 10 μl of DNA construct was passed through the channel and incubated for 5 minutes for the biotin-labeled DNA constructs to bind the streptavidin on the glass surface. The chamber was then washed with PBS + 0.1% Tween 20 to remove unbound constructs. DNA coated microspheres are passed into the chamber and the chamber was sealed with vacuum grease, incubted for 2 minutes for the complementary strands to hybridize and then loaded into the CFM just before the experiment was performed.

### Experimental protocol

To set up for an experiment, the camera needs to be plugged into the router and the battery connected to power the CFM. Once these connections are made, the Wi-Fi signal from the router will appear in the network option in the computer. Once the connection from the router is established, the camera can be accessed from the LabVIEW™ program (Figure S7). The run button on the LabVIEW™ toolbar activates the streaming from the camera. The sealed chamber was screwed into the base of the lens tube (Thorlabs, Part # SM1L05), and the lens tube was then screwed into the rest of the optical assembly until the beads were in focus. The CFM was then loaded into the centrifuge bucket and the speed was controlled from the front panel of the centrifuge. The force generated on the tether is the centrifugal force experienced by the beads F = mω^2^r, where m is the effective mass of the bead (actual mass-mass of buffer displaced), ω is the angular velocity and r is the distance from the center of the rotar to the chamber (0.133 m). The effective mass of beads were determined to be 6.9*10^-12^g for the Dynabeads™ M-270 (www.thermofisher.com) by previous report (5) and 2.6*10^-12^ g for 5.2 μm polystyrene beads (Catalogue # SVP-50-5, www.spherotech.com). Mass calculation for the beads and the RPM used to achieve the force used in the experiments are shown in Table S4 and Table S5.

### Data analysis

The data obtained from the CFM was analysed using custom written MATLAB^®^ (2019) program. The program identifies beads using inbuilt “imfindcircles” algorithm with a user override for non-spherical beads and dirt wrongly identified as beads. Once the beads are identified from the image at the start of the experiment, the software calculates the variance of the image intensity at the bead location for all the frames. When beads dissociate, it is indicated by the sharp drop in variance. Multiple drops in variance observed are due to break in multiple-tethered beads, which are excluded from the analysis. The MATLAB code for this program is provided in supplementray materials (File S8). The data plots are constructed in OriginPro 2018 and fitting of data was done using inbuilt single exponential decay function, y = y_0_ + A*exp (-kt), where y is the fraction of tethers remaining at a given time t, y_0_ is the y-axis offset or the baseline, A is the fraction of tethers at the beginning of the experiment (typically 1) and k is the off-rate for that particular force. Off-rate for any given condition is determined by at least triplicate experiments where individual k values were determined separately, and data reported as the mean and standard deviation of the replicates.

## Results

We had two main improvement goals in the development of our newest CFM: to establish real-time wireless control and streaming, and to ensure an upgrade pathway for future CFM generations. Our previous CFM setup using the Intel Edison computer required running “blind”, with images stored on limited onboard memory and later transmitted wirelessly for analysis, providing a frustrating user experience. Furthermore, the Edison computer required programming skills to implement changes, and its abrupt discontinuation by Intel made us realize how difficult simple upgrades could be. To solve these challenges, we implemented a Wi-Fi streaming strategy using an onboard router coupled with a GigE (gigabit Ethernet) vision camera. Both Wi-Fi and GigE vision follow standardized protocols that are routinely updated as technology progresses, providing a measure of future proofing for our CFM as wireless and camera technologies continue to improve. By using these standard communication protocols, we are also free to use any common programming language (including LabVIEW™) for instrument control.

To implement this strategy, we started from the basic microscopy module developed previously (8), and substituted the USB camera from that design to a gigabit Ethernet camera. We next identified two differerent Wi-Fi routers that were small enough to fit within our microscopy module inside a centrifuge bucket. The first was a commercial travel router (TP-Link AC750) with a theoretical bandwidth of 100 Mbps limited by the Ethernet port speed, and the second was an industrial grade OEM router (EmbedAir1000 from Acksys Communications and Systems) with a theoretical bandwidth of 557 Mbps limited by the wireless radio capabilities. We devised a networking scheme to establish a virtual ethernet connection between the camera and the computer by connecting the camera to the router in access point mode and accessing the Wi-Fi signal from a desktop computer (Figures 2a, 2b). A step-by-step networking guide for both routers are provided in the supplementary notes 2 and 3.

**Figure 2:**
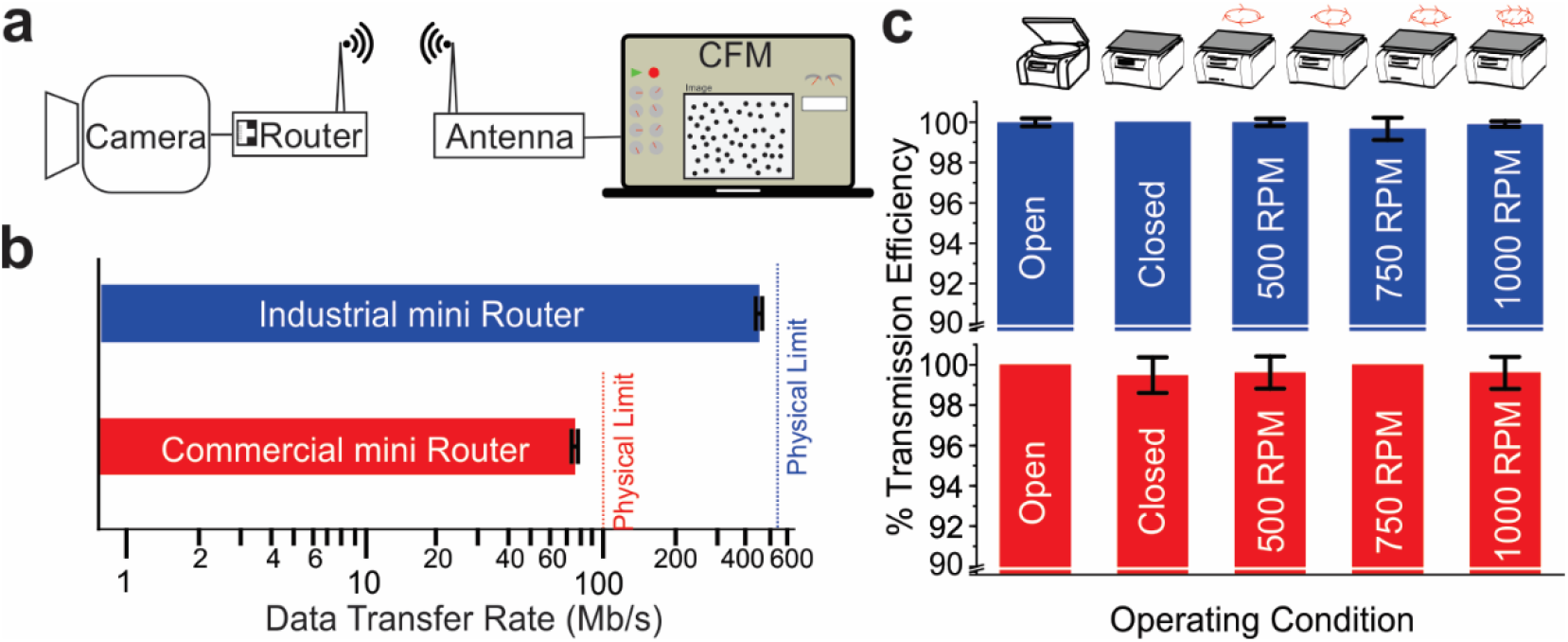
CFM networking strategy and streaming quality. (a) Schematic diagram showing wireless communication between the camera and the computer terminal. (b) Measured data transfer rates using a commercial mini router and an industrial grade OEM router. (c) Quantified data transfer quality in various conditions using commercial mini router (red bars) and industrial grade OEM router (blue bars) shows that the data transfer is >99% reliable. Very few images have lost data packets which appear as dark pixels.

Once the basic connection scheme was established, we designed new 3D printed housings to hold the CFM components including the new routers and support electronics. We also developed a wiring strategy to ensure the router and the camera have consistent power to remain connected for long periods in the centrifuge. With the router operating at 5 V and the GigE camera operating at 12-24 V, we used separate 5 V and 12 V voltage step-up converters wired in series to a Lithium ion battery (Wiring diagram in Figure S2, S4). We powered the LED light source separately from a coin cell battery. Using this configuration with a 2000 mAh battery resulted in a run time exceeding 2 hours. We developed a new LabVIEW™ program to control the camera, to display and store images, and to adjust relevant parameters such as frame rate, area of interest and exposure time (Figure S7, Supplementary file 7). Using this program, the captured images and the associated meta data including timestamp are transferred to the computer in real time, displayed on screen, and stored on the computer hard disk for analysis.

For our first tests of the new CFM, we assessed data transmission with each of the routers. We were able to consistently transmit data at a rate of 77.059 ± 0.002 Mbps with the commercial travel router and 454.3 ± 0.6 Mbps data with industrial router (Figure 2b). These rates equate to 1.9 frames per second (fps) and 11.3 fps in full frame mode for our 5 MP camera, and both represent ~80% of the theoretical bandwidth of their respective routers. Faster frame rates can be obtained by reducing the frame sizes, but the maximum frame rates are often set by camera firmware. Next we assessed transmission efficiency, since wireless transmission can suffer from dropped packets. The GigE vision standard uses UDP protocol which has a higher data transfer rate than the common TCP protocol but does not guarantee packet delivery. We analyzed the data stream in different scenarios: (i) Stationary CFM inside centrifuge with opened lid, (ii) stationary CFM inside the centrifuge with closed lid, (iii) spinning CFM with closed lid at 500, 750, and 1000 rpm. We found that data transmission was not affected by closing the lid or spinning the device in the centrifuge, despite the thick walls of the centrifuge. We observed that both routers were able to transmit data with >99% reliability (Figure 2c) and packet loss appeared random.

Having validated the technical performance of our instrument’s operation, we next developed a MATLAB^®^ post-processing program for the analysis of single molecule experiments from the CFM (supplementary file 8). For a model system, we considered a generic off-rate experiment in which microspheres tethered to a cover glass are exposed to force and dissociate over time (Figure 3a). We implemented this experiment using two biotinylated DNA handles that attach to either streptavidin coated beads or cover glass and form tethers when complementary single-stranded overhangs bind to each other (Figure 3a). The DNA tethered beads were observed in the CFM module and subjected to centrifugal force, with DNA shearing observed by bead detachment from the cover glass. The program was designed to identify tethered beads for analysis with a circle finding algorithm and display color coded circles on the images (Figure 3b-d). After identifying tethered beads, the software scanned all frames (Figure 3e) and constructed time traces of each bead to visually identify rupture frames (Figure 3f). For each bead in each frame, the image variance was calculated for a region of interest and was used to indicate rupture events (Figure 3g), which were plotted to display decay kinetics (Figure 3g, inset).

**Figure 3:**
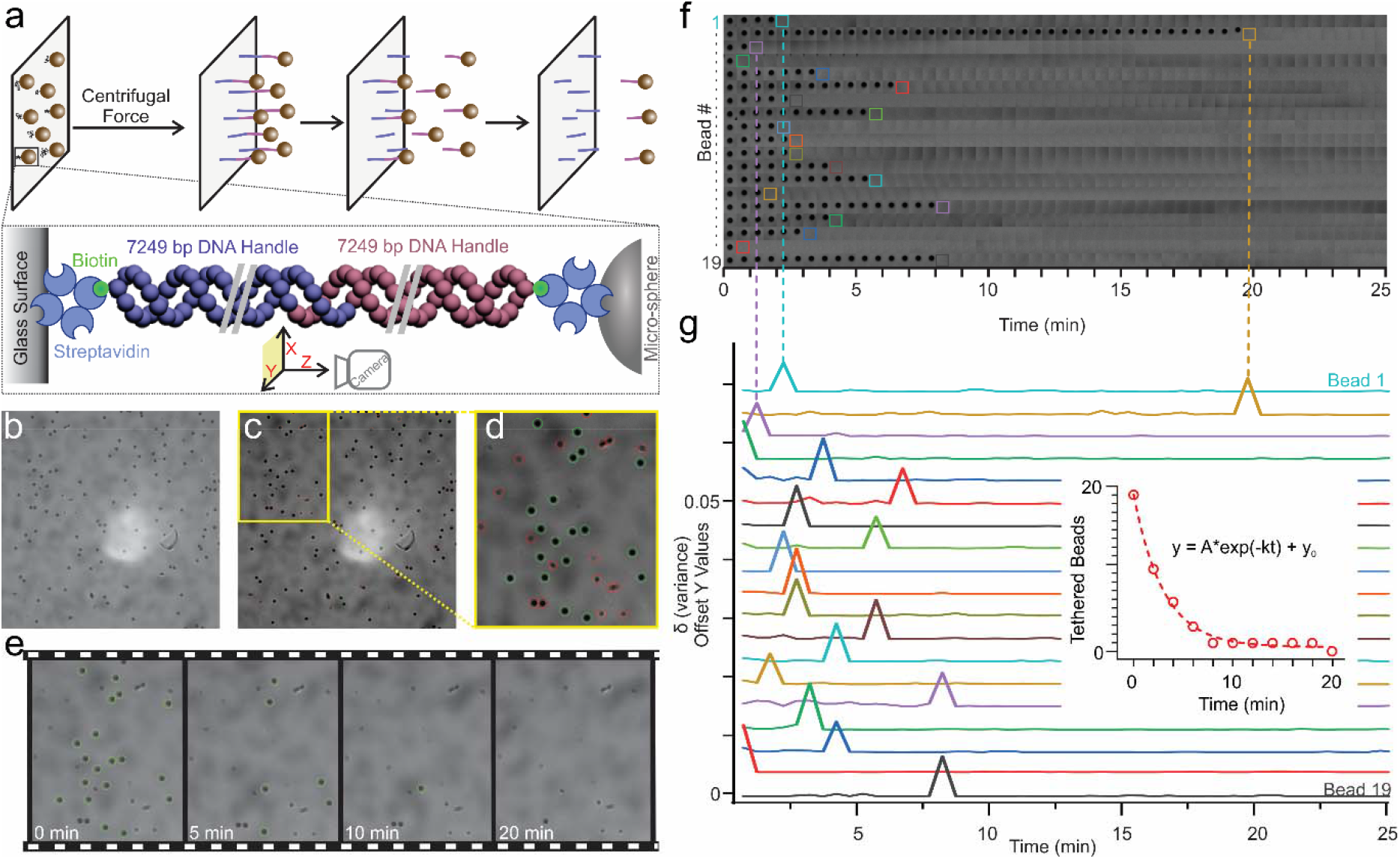
Experiment schematic and data analysis. (a) Schematic of single-molecule experiment. The DNA handles with the sequence of interest is tagged to a glass side and a microbead. The hybridized DNA construct will unbind when force is exerted while spinning the CFM. (b) Image displaying tethered beads under tenstion through the axis of view of the camera. (c) Image showing bead selected through the MATLAB^^®^^ program for further analysis (green) and excluded beads (red). (d) Expanded view of a section of previous image (e) Images of area of interestat different time points. It can be observed that the beads disappear as time progresses due to tether rupture under force. (f) Tracking of individual beads over time. (g) Derivative plot of variance in the pixel intensity around the bead. The variance will change when the bead dissociates from the surface, and this will appear in derivative plot as a peak. (Inset) Cumulative decay plot of remaining tethers at a given point of time (red circles) is constructed from the derivative plot and kinetic parameters such as off-rates at a particular force can be obtained by fitting the cumulative with a single exponential decay function (red line).

To demonstrate the use of the new CFM and software for single-molecule experiments, we performed DNA shearing experiments that showcase the dependence of force, length, and GC content in dissociation kinetics. Using the pair of cross-hybridizing biotinylated DNA handles described above, we probed the kinetics of incorporation of different sequences in the short duplex (Figures 4a, S8). First we investigated force-dependent dissociation of a 7 bp interaction using force clamp experiments at 2-12 pN (Figure 4b). The off-rates were determined by fitting the cumulative decay rates with a single-exponential decay curve, and ranged from 0.0034 ± 0.0009 s^-1^ to 0.0015 ± 0.0001 s^-1^. We extrapolated the thermal off-rate by fitting this data with Bell-Evans model (Figure 4c) (11, 12), where the force dependent dissociation rate is described as k_off_ (f) = k_off_ exp (f/f_β_), where k_off_ (f) is the off-rate at the applied force f, k_off_ is the thermal off-rate, and f_β_ is the characteristic force scale. Using this model, we found a thermal off-rate k_off_ of 0.0013 ± 0.0001 s^-1^ and a force scale f_β_ of 13.0 ± 1.6 pN.

**Figure 4:**
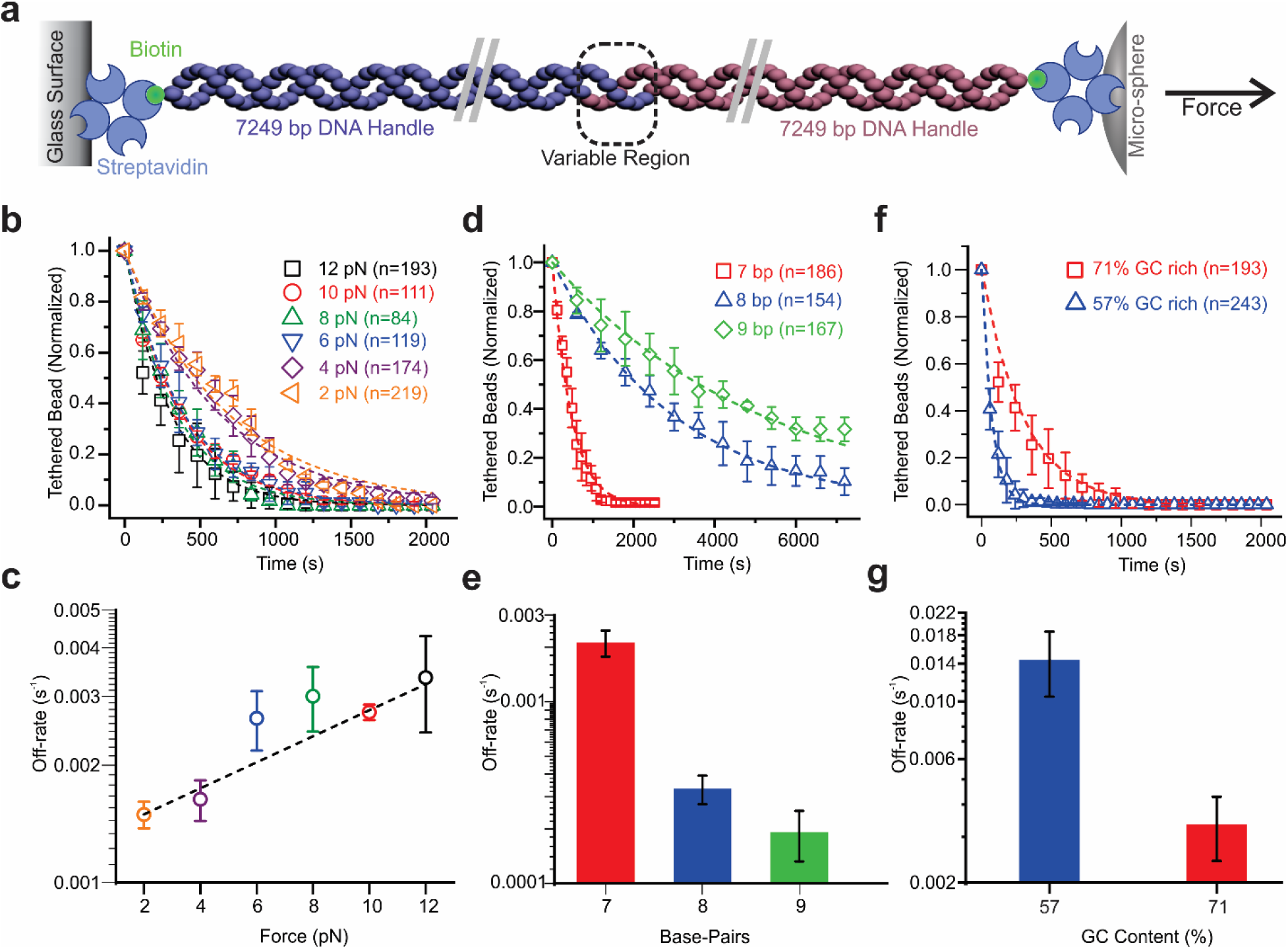
Single-molecule force clamp experiments with the CFM. (a) Schematics of experiment where force is applied to analyze the strength of DNA structures hybridized through complementary overhangs. (b) Decay plot showing the effect of force (2 pN to 12 pN) for a 7 bp dissociation. Off-rates of the molecules can be determined by fitting the data with a single exponential decay (dotted lines). (c) Off-rates determined at various forces can be used to determine thermal off-rates by fitting the data with Bell-Evans model. (d) Decay plot showing the dependency of length of the overhang on force induced dissociation when held at a constant force of 4.5 pN. (e) Off-rates of DNA shearing experiments containing 7 bp - 9 bp at 4.5 pN. (f) Decay plot showing the dependency of GC content on DNA shearing when held at a constant force of 12 pN. (g) Off-rates of DNA shearing experiments containg 7 bp at 12 pN, but varied GC content by single GC to AT mutation.

We also performed force clamp experiments at 4.5 ± 1.3 pN on DNA duplexes with lengths of 7 bp, 8bp, and 9 bp to study the influence of DNA length and its thermodynamic stability (Figure 4d). We found the length dependent off-rates to be 0.0021 ± 0.0003 s^−1^, 0.00032 ± 0.00006 s^−1^, 0.00027 ± 0.00006 s^−1^, respectively for 7, 8, and 9 bp (Figure 4e). This remarkable difference in the force-induced off-rates suggests that this approach would also be sensitive to the GC content of the sequences, thereby opening up platforms for single-nucleotide polymorphism (SNP) detection and force fingerprinting. To test the sensitivity of this approach, we designed an additional DNA construct based on the 7 bp interaction that replaced a single G-C pair with an A-T pair. We subjected both variants to a 12± 1.4 pN force clamp and measured the dissociation rates (Figure 4f) to be 0.0034 ± 0.0009s^-1^ for the native duplex and 0.0145 ± 0.004 s^-1^ for the weakened duplex (Figure 4g). This indicates that the method is sensitive enough to distinguish a single-nucleotide variation in DNA fragments.

## Discussion

In this work, we have unveiled a next generation CFM that is completely plug-and-play with high-bandwidth live streaming over Wi-Fi. This instrument is the first CFM that offers similar performance to the fiber-optic wired CFMs (5, 6), but with the convenience and simplicity of a plug-and-play module (8). Our design also for the first time considers future upgrades and ease of programming, opting for commercially available Wi-Fi components that are interchangeable and have standardized protocols. These design changes are centered around the user experience, and in making single-molecule experiments easier to perform. In that regard, the CFM is uniquely well suited among single molecule techniques, and can even be used by undergraduate students and researchers without a technical biophysics training.

As with previous builds, the CFM has orders of magnitude price difference compared with most single-molecule setups. The CFM module itself can be constructed inexpensively with choice of camera and objective, and most labs already have access to benchtop centrifuges that can be used. In the supplemental information, we have provided eall the details necessary to reconstruct the instrument including 3D models, parts lists, and step-by-step protocols.

Since the initial development of the CFM, there have been several other techniques that were developed to provide multiplexed single-molecule experimentation. In particular, there have been significant advances in multiplexed magnetic tweezers (13, 14) as well as development of new technologies like acoustic force spectroscopy (AFS) (15) and optical pushing (16). These methods have helped expand the field, especially with commercialization of AFS, but we still believe that the CFM offers some distinct features for ease of use, low cost, and a wide and calibration-free force range. With this current development, we further advance the state-of-the-art in CFM design and make it easier than ever to build and use.

## Acknowledgements

Research reported in this publication was supported by the NIH through NIGMS under award R35GM124720 to K.H. We thank Dr. Tony Hoang for technical help and suggestions, the UAlbany Research IT team for help with networking, and Dr. Arun Richard Chandrasekaran for discussions on the project, and for critical suggestions and editing on the manuscript.

## References

1. Ritort, F. 2006. Single-molecule experiments in biological physics: methods and applications. J Phys Condens Matter. 18:R531–583.

2. Deniz, A.A., S. Mukhopadhyay, and E.A. Lemke. 2008. Single-molecule biophysics: at the interface of biology, physics and chemistry. Journal of The Royal Society Interface. 5:15–45.

3. Zlatanova, J., and K. van Holde. 2006. Single-Molecule Biology: What Is It and How Does It Work? Molecular Cell. 24:317–329.

4. Nathwani, B., W.M. Shih, and W.P. Wong. 2018. Force Spectroscopy and Beyond: Innovations and Opportunities. Biophysical Journal. 115:2279–2285.

5. Halvorsen, K., and W.P. Wong. 2010. Massively Parallel Single-Molecule Manipulation Using Centrifugal Force. Biophysical Journal. 98:L53–L55.

6. Yang, D., A. Ward, K. Halvorsen, and W.P. Wong. 2016. Multiplexed single-molecule force spectroscopy using a centrifuge. Nature Communications. 7:11026.

7. Yang, D., and W.P. Wong. 2018. Repurposing a Benchtop Centrifuge for High-Throughput Single-Molecule Force Spectroscopy. In: Peterman EJG, editor. Single Molecule Analysis: Methods and Protocols. New York, NY: Springer. pp. 353–366.

8. Hoang, T., D.S. Patel, and K. Halvorsen. 2016. A wireless centrifuge force microscope (CFM) enables multiplexed single-molecule experiments in a commercial centrifuge. Review of Scientific Instruments. 87:083705.

9. Kirkness, M.W.H., and N.R. Forde. 2018. Single-Molecule Assay for Proteolytic Susceptibility: Force-Induced Collagen Destabilization. Biophysical Journal. 114:570–576.

10. Jin, L., L. Kou, Y. Zeng, C. Hu, and X. Hu. 2019. Sample preparation method to improve the efficiency of high-throughput single-molecule force spectroscopy. Biophys Rep. 5:176–183.

11. Bell, G.I. 1978. Models for the specific adhesion of cells to cells. Science. 200:618–627.

12. Evans, E., and K. Ritchie. 1997. Dynamic strength of molecular adhesion bonds. Biophysical Journal. 72:1541–1555.

13. De Vlaminck, I., and C. Dekker. 2012. Recent Advances in Magnetic Tweezers. Annual Review of Biophysics. 41:453–472.

14. Vlaminck, I.D., T. Henighan, M.T.J. van Loenhout, D.R. Burnham, and C. Dekker. 2012. Magnetic Forces and DNA Mechanics in Multiplexed Magnetic Tweezers. PLOS ONE. 7:e41432.

15. Sitters, G., D. Kamsma, G. Thalhammer, M. Ritsch-Marte, E.J.G. Peterman, and G.J.L. Wuite. 2015. Acoustic force spectroscopy. Nature Methods. 12:47–50.

16. Sitters, G., N. Laurens, E. de Rijk, H. Kress, E.J.G. Peterman, and G.J.L. Wuite. 2016. Optical Pushing: A Tool for Parallelized Biomolecule Manipulation. Biophysical Journal. 110:44–50.

