## Supplementary Information for "A Wi-Fi live streaming Centrifuge Force Microscope for benchtop single-molecule experiments"

**Contents**

Figure S1: Assembling the optical components of the CFM.

Figure S2: Wiring diagram for CFM with the mini-router with list of all components.

Figure S3: Sketch of fully assembled CFM from various angles.

Figure S4: Wiring diagram for CFM with the OEM router with list of all components.

Figure S5: Sketch of fully assembled CFM from various angles.

Figure S6: Sketch of counter balance for the CFM.

Figure S7: LabVIEW front panel view.

Figure S8: DNA construct preparation.

Supplementary Note 1. Instructions for counterbalance design.

Supplementary Note 2. Step-by-step networking instructions for mini router (TP-link).

Supplementary Note 3. Step-by-step networking instructions for OEM router.

Table S1. List of components.

Table S2. Resend Parameters.

Table S3. List of oligonucleotide sequences.

Table S4: Mass calculation for micro-spheres.

Table S5: RPM used to achieve the prescribed force for M270 and SVP-50-5 beads.

**Other Supplementary Files**

1. 3D files for housing – Mini Router
   a. CFM Housing – Fusion Drawing

b, c. STL file

2. 3D files for housing – OEM Router Housing (Optics housing is the same as the mini-router)
   a. Fusion Drawing
   b. STL file
3. 3D files for counterbalance
   a. Mini Router – Fusion Drawing
   b. STL file
4. 3D Files for the counterbalance
   a. OEM Router – Fusion Drawing
   b. STL file
5. Custom print connector for Blackfly camera
   a. Fusion Drawing
   b. STL file
6. Custom print pinhole 4mm diameter
   a. Fusion Drawing
   b. STL file
7. LabVIEW^TM^ file
8. Matlab® file

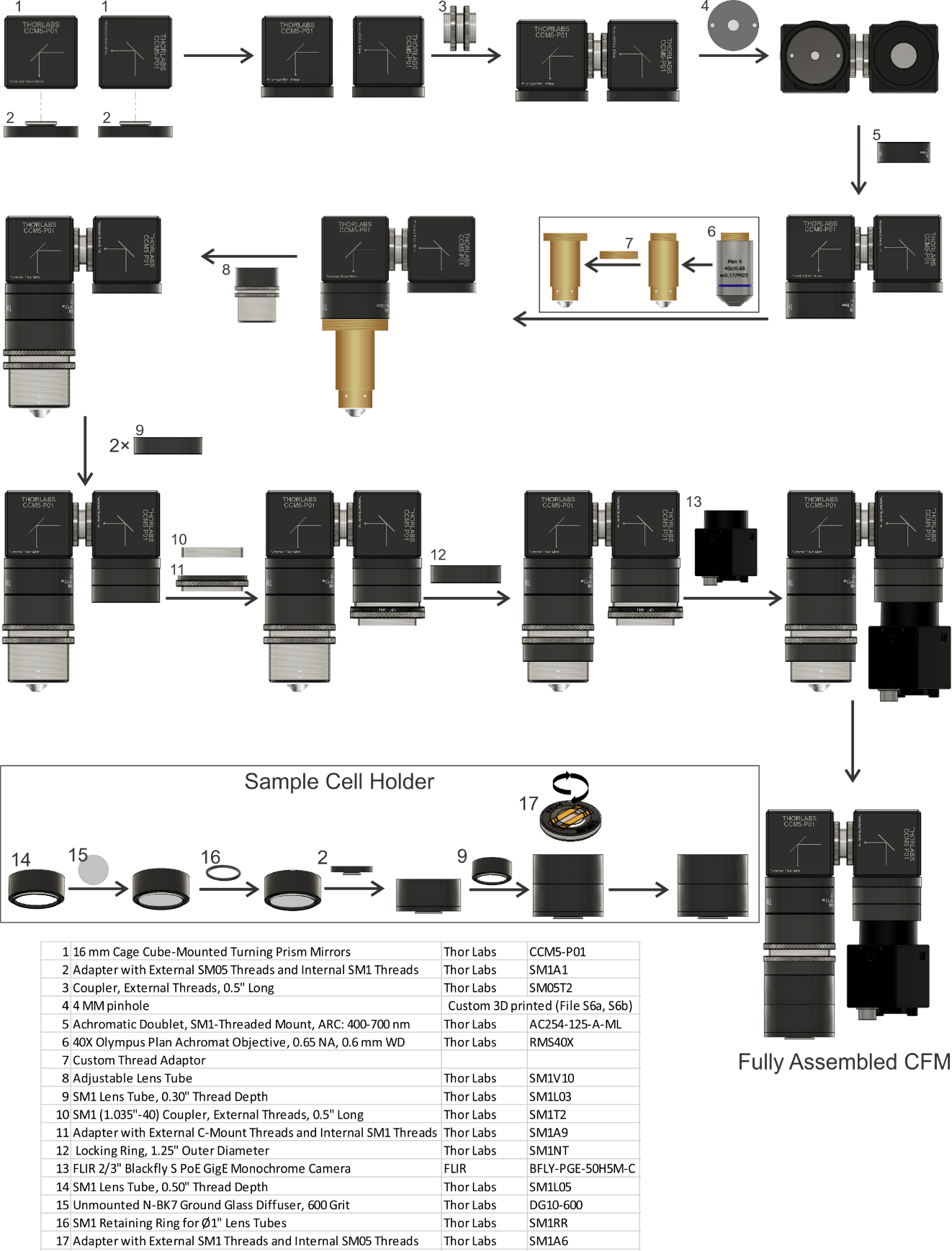

**Figure S1**: Assembling the optical components of the CFM.

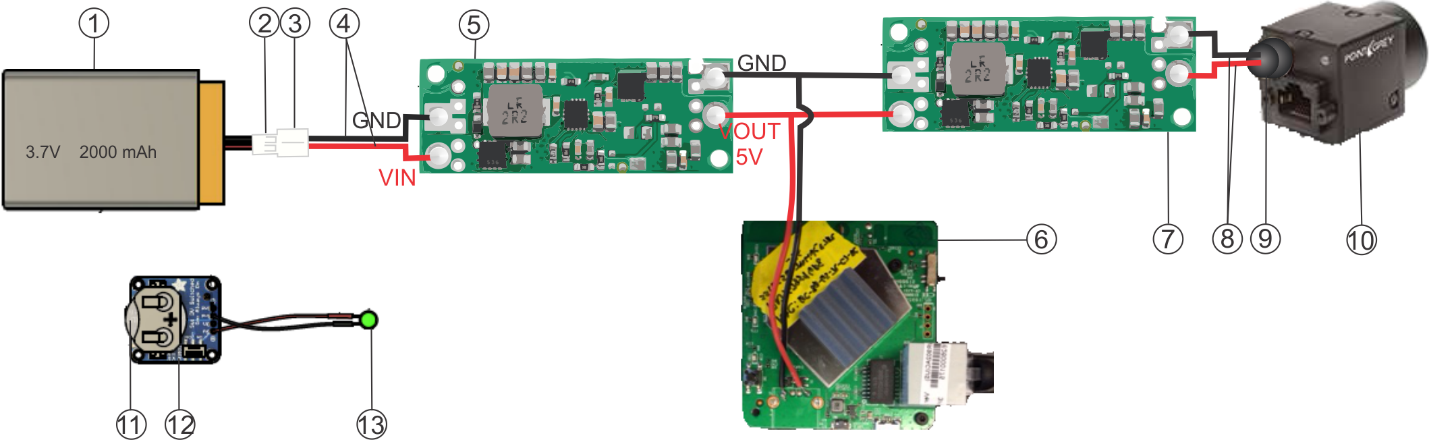

| **Item #** | **Name** | **Source** | **Part/Model Number** |
| --- | --- | --- | --- |
| 1 | Lithium Ion Battery - 3.7v 2000mAh | www.adafruit.com | 2011 |
| 2 | JST PH 2-Pin Cable - Female Connector | www.adafruit.com | 261 |
| 3 | JST PH 2-Pin Cable – Male Header | www.adafruit.com | 3814 |
| 4 | Silicone Cover Stranded-Core Wire Red & Black | www.adafruit.com | 3165 & 3164 |
| 5 | 5V Step-Up Voltage Regulator U3V70F5 | www.pololu.com | 2891 |
| 6 | TP-Link AC750 Wireless Portable Nano Travel Router | www.amazon.com | TL-WR902AC |
| 7 | 12V Step-Up Voltage Regulator U3V70F12 | www.pololu.com | 2895 |
| 8 | female jumper wires | www.adafruit.com | 1919 |
| 9 | Custom print 8 pin GPIO connector housing female jumper wires | Supplementary file S5a, 5b | |
| 10 | Gigabit Ethernet camera | www.flir.com | BFLY-PGE-50H5M-C |
| 11 | CR2032 Lithium Coin Cell Battery | www.adafruit.com | 654 |
| 12 | 12mm Coin Cell Breakout w/ On-Off Switch | www.adafruit.com | 1867 |
| 13 | Green (525 nm) LED | www.thorlabs.com | LED525E |

**Figure S2**: Wiring diagram for CFM with the mini-router with list of all components.

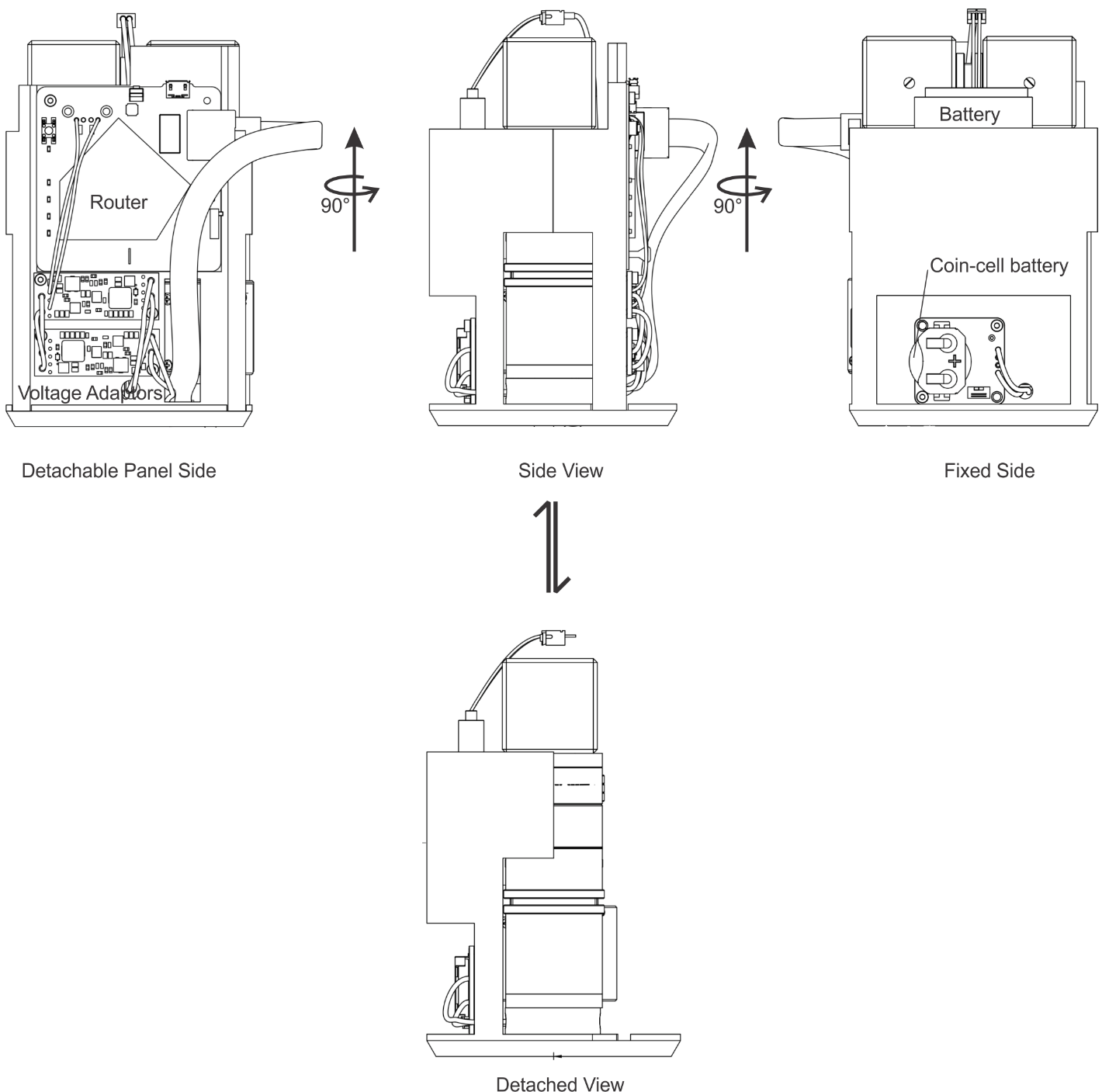

**Figure S3**: Sketch of fully assembled CFM from various angles. The electronics panel detach from the main housing which consist of base and the battery holder. The 3d files for printing the housing are provided in supplementary files S1a, 1b, 1c. The optical components can be assembled in this open configuration, before fully assembling the unit and loading it into the CFM.

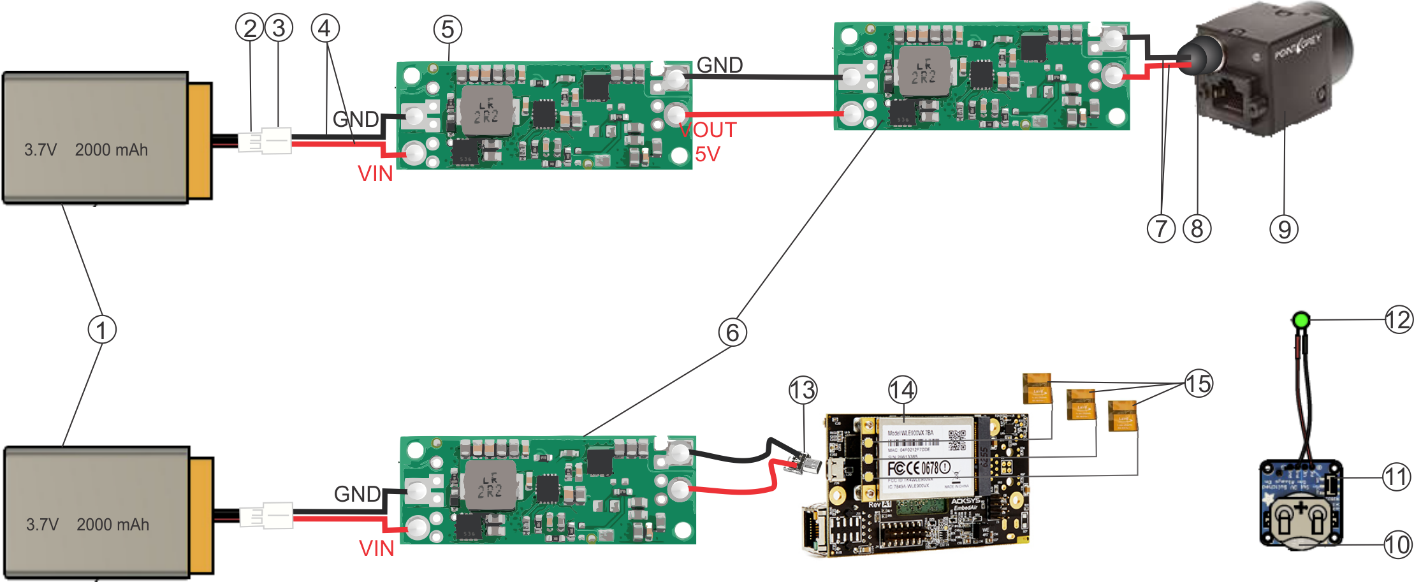

| **Item #** | **Name** | **Source** | **Part/Model Number** |
| --- | --- | --- | --- |
| 1 | Lithium Ion Battery - 3.7v 2000mAh | www.adafruit.com | 2011 |
| 2 | JST PH 2-Pin Cable - Female Connector | www.adafruit.com | 261 |
| 3 | JST PH 2-Pin Cable – Male Header | www.adafruit.com | 3814 |
| 4 | Silicone Cover Stranded-Core Wire Red & Black | www.adafruit.com | 3165 & 3164 |
| 5 | 5V Step-Up Voltage Regulator U3V70F5 | www.pololu.com | 2891 |
| 12 | 12V Step-Up Voltage Regulator U3V70F12 | www.pololu.com | 2895 |
| 7 | female jumper wires | www.adafruit.com | 1919 |
| 8 | Custom print 8 pin GPIO connector housing female jumper wires | Supplementary file S5a, 5b | |
| 9 | Gigabit Ethernet camera | www.flir.com | BFLY-PGE-50H5M-C |
| 10 | CR2032 Lithium Coin Cell Battery | www.adafruit.com | 654 |
| 11 | 12mm Coin Cell Breakout w/ On-Off Switch | www.adafruit.com | 1867 |
| 12 | Green (525 nm) LED | www.thorlabs.com | LED525E |
| 13 | Right Angle Micro B Plug Down | www.adafruit.com | 4105 |
| 14 | EmbedAir1000 Wi-Fi router | www.acksys.com | EmbedAir1000 |
| 15 | Antennas Dual-Band mFlexPIFA, 120mm, MHF1 | www.mouser.com | 239-001-0034 |

**Figure S4**: Wiring diagram for CFM with the OEM router with list of all components. Here, unlike the mini-router, the router and battery are in separate circuits and has separate housings and should be loaded into different centrifuge buckets. Detailed sketch is given in figure S5.

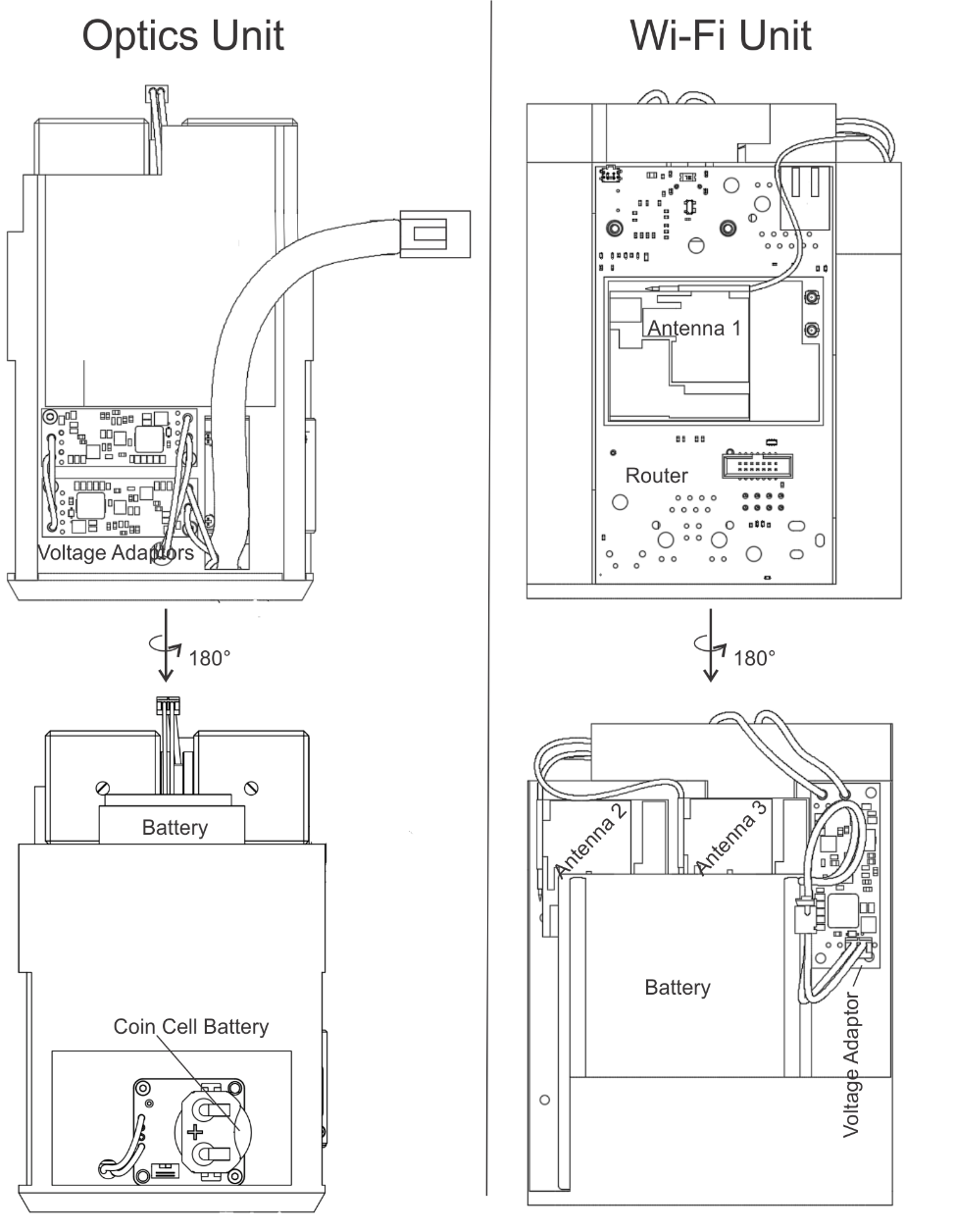

**Figure S5**: Sketch of fully assembled CFM from various angles. The electronics panel detach from the main housing which consist of base and the battery holder. The 3D files for printing the housing are provided in supplementary files S1a, 1b, 1c, 2a, 2b. The optical components can be assembled in this open configuration, before fully assembling the unit and loading it into the CFM.

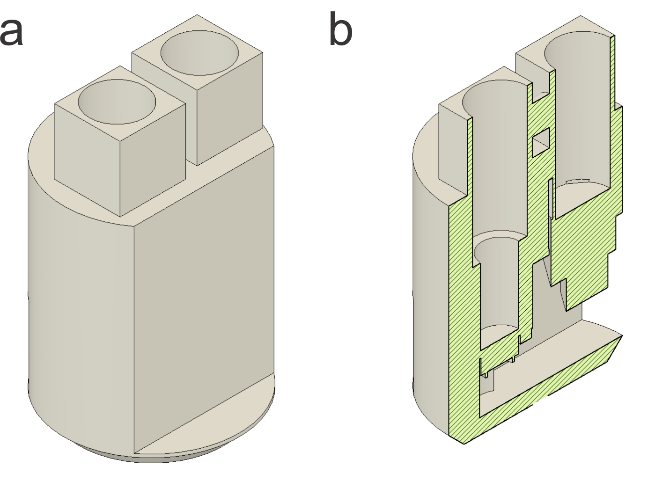

**Figure S6**: Sketch of counter balance for the CFM. (a) Intact counterbalance for the mini-router setup. Quarters can be added to the wells to match the mass of CFM. The 3D files for printing the counterbalances for the mini-router and the OEM router are given in supplementary files S3a, 3b, 3c and 3d. (b) Cutaway view of the counterbalance.

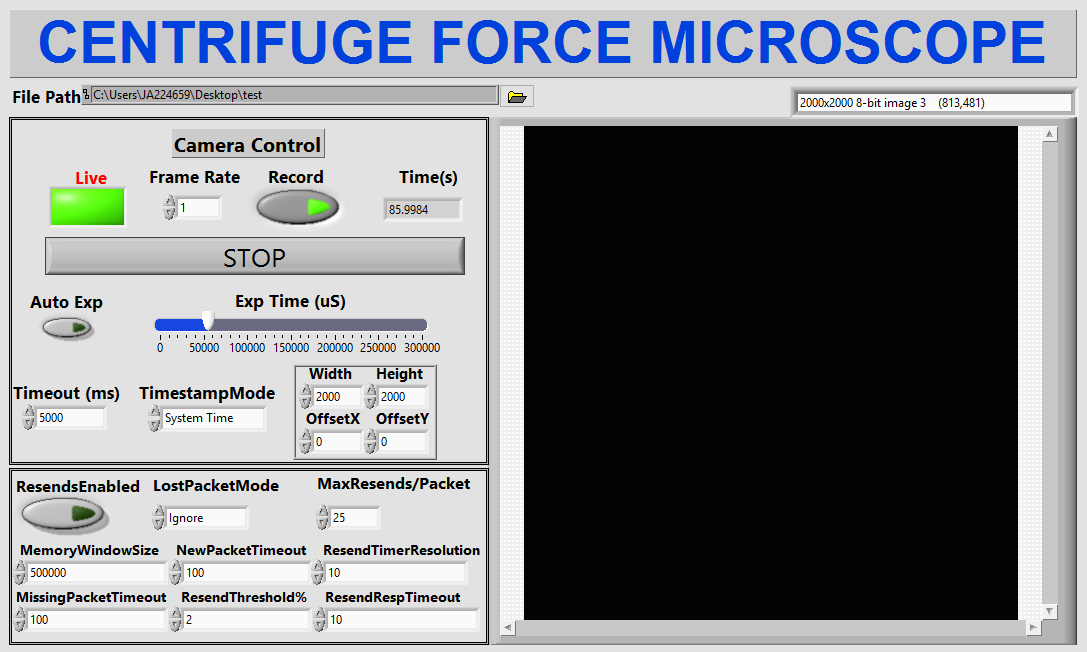

**Figure S7**: LabVIEW front panel view. This program can be used to remotely access and control the camera from the computer and stream data in real time. Data will be stored in the hard disk.

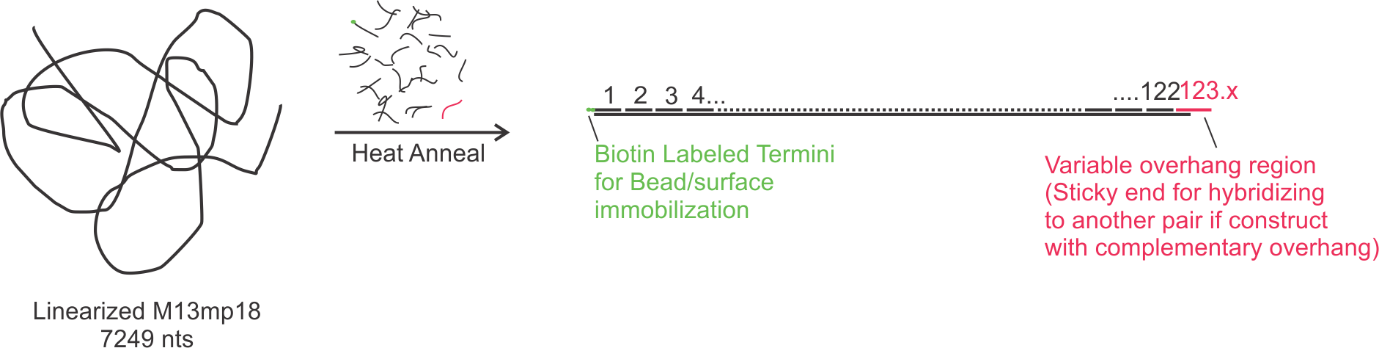

**Figure S8**: DNA construct preparation. Linearized M13mp18 single-stranded DNA is annealed with the backbone oligos, the 3ˊ overhang oligo (red strand) and terminal oligo labelled with biotin (green label). The biotin is utilized to immobilize the DNA construct to a streptavidin coated glass surface or microsphere. The overhang is used for hybridization to a similar DNA construct with complementary overhang sequence for the single-molecule pulling experiment.

Supplementary Note 1. Instructions for counterbalance design.

The CFM’s counterbalance is a 3D printed geometric replica of the instrument, including its housing with two 1-inch holes for adjusting the center of mass.

1. To make the CFM mock up, measure the dimensions of the CFM with a caliper and use these measurements to model the instrument in a CAD program. Alternatively, prepare a 3D model of the instrument using .step files from the manufacturers’ website, and model a body that occupies the same volume as the model CFM.
2. Drill two 25.25 mm holes centered on the cylindrical portions of the CFM to a depth of 60 mm. Save the model.
3. Import the model of the CFM housing into the same file as the CFM mock up.
4. Combine both halves of the CFM housing with the mock up CFM. This completes counterbalance; however, it must be mirrored across the X axis to position its center of mass exactly parallel to the instrument’s.
5. Create a mirror image of the counterbalance by selecting the component and choose the mirror tool and then select the X axis as the mirror plane. Complete the operation.
6. Create an .stl file of the counterbalance model and import it into a slicing program such as Cura 4.0.
7. Slice the counterbalance in Cura or a similar program using the same settings as for the housing, but with these changes:
   - Layer Height = 0.4 mm
   - Infill Density = 40%
8. After printing the counterbalance, add quarters to match the mass of your CFM.
9. It is very important to match the center of mass of CFM and the counter balance to minimize the vibration while running. This has to be determined through trial and error by by placing quarters at various height levels, using 3D printed hollow cylinder of various heights.

Supplementary Note 2. Step-by-step networking instructions for mini router (TP-link).

1. Plug in wireless router and power on, then connect to PC.
2. Open “Network Connections” on PC and choose “TP-Link_XXXX”, where XXXX designates your specific router.
3. Open a web browser and type tplinkwifi.net. At the login screen type “admin” for Login and Password 🡪 Enter (“admin” is the default user id and password for tplink routers).
4. Under Operation Mode tab select “Access Point”.
5. Set IP to “Static IP” with the following settings:

IP Address: 192.168.0.1 Subnet Mask: 255.255.0.0

Click “Save”.

6. Go to Windows “Control Panel” 🡪 Network and Internet 🡪 Network and Sharing Center 🡪 Change Adapter Settings (left panel) 🡪 Choose “Ethernet” and “Wi-Fi” connections icons simultaneously and choose “Bridge Connections” from the right-click dropdown list. If you are connected to internet using the Ethernet, you may want to unplug the connection for smooth connectivity to the router. If you have two Ethernet ports, choose the one that is not connected to the internet to bridge to your Wi-Fi connection from TP-Link router.
7. Now the camera will be discoverable in the NI MAX or the camera SDK. If the camera is not discovered in the NI MAX, you can force IP configuration using your camera SDK where the router and the camera will share the same IP and thereby establishing virtual Ethernet connection between camera and the computer.

Supplementary Note 3. Step-by-step networking instructions for OEM router.

1. Power the Acksys router, and Open “Network Connections” on PC and choose “Acksys”
2. Using your web browser, open the WEB page at address 169.254.74.253 (you may have to experimentally figure out based on your network), and select the SETUP tab. The main page is the PHYSICAL INTERFACES OVERVIEW. First, select the country and click the Save button.
3. You must now click on the “Edit” button for the Wi-Fi Connection under the “WIFI Interface”, under “ACTIONS”:
4. In the WIRELESS SETTINGS page,
   a. Check the “Enable device” box.
   b. Select the “802.11a+n 5GHz” as the “802.11 mode”.
   c. Select 40 or 800 MHz in “HT mode”.
   d. Uncheck “Automatic channel select”, and select channel 36 and 149.
5. In the same page, under “Interface configuration”
   a. Choose “Role” as “Access Point”.
6. Save and Apply.
7. Go to Windows “Control Panel” 🡪 Network and Internet 🡪 Network and Sharing Center 🡪 Change Adapter Settings (left panel) 🡪 Choose “Ethernet” and “Wi-Fi” connections icons simultaneously and choose “Bridge Connections” from the right-click dropdown list. If you are connected to internet using the Ethernet, you may want to unplug the connection for smooth connectivity to the router. If you have two Ethernet ports, choose the one that is not connected to the internet to bridge to your Wi-Fi connection from Acksys router.
8. Now the camera will be discoverable in the NI MAX or the camera SDK. If the camera is not discovered in the NI MAX, you can force IP configuration using your camera SDK where the router and the camera will share the same IP, thereby establishing virtual Ethernet connection between camera and the computer.

**Table S1.** List of components.

| **Component** | **Vendor** | **Part No.** | **Price** | **Quantity** |
| --- | --- | --- | --- | --- |
| FLIR 2/3" Blackfly S PoE GigE Monochrome Camera (BFLY-PGE-50H5M-C) | [FLIR](https://www.flir.com/products/blackfly-gige/?model=BFLY-PGE-50H5M-C) | BFLY-PGE-50H5M-C | $995.00 | 1 |
| THORLABS SM1L03 - SM1 Lens Tube, 0.30"Thread Depth | [Thorlabs](https://www.thorlabs.com/thorproduct.cfm?partnumber=SM1L03) | SM1L03 | $12.16 | 2 |
| THORLABS SM1A1 Adapter with External SM05 Threads and Internal SM1 Threads | [Thorlabs](https://www.thorlabs.com/thorproduct.cfm?partnumber=SM1A1) | SM1A1 | $21.22 | 3 |
| THORLABS CCM5-P0116 mm Cage Cube-Mounted Turning Prism Mirrors | [Thorlabs](https://www.thorlabs.com/thorproduct.cfm?partnumber=CCM5-P01) | CCM5-P01 | $139.73 | 2 |
| THORLABS SM05T2 SM05 (0.535"-40) Coupler, External Threads, 0.5" Long | [Thorlabs](https://www.thorlabs.com/thorproduct.cfm?partnumber=SM05T2) | SM05T2 | $19.33 | 1 |
| RMS40X - 40X Olympus Plan Achromat Objective, 0.65 NA, 0.6 mm WD | [Thorlabs](https://www.thorlabs.com/thorproduct.cfm?partnumber=RMS40X) | RMSX40 | $680.78 | 1 |
| THORLABS SM1L05E - SM1 Lens Tube Without External Threads, Optic Retention Lip, 1/2" Thread Depth | [Thorlabs](https://www.thorlabs.com/thorproduct.cfm?partnumber=SM1L05E) | SM1L05E | $12.93 | 1 |
| THORLABS SM1NT - SM1 (1.035"-40) Locking Ring, 1.25" Outer Diameter | [Thorlabs](https://www.thorlabs.com/thorproduct.cfm?partnumber=SM1NT) | SM1NT | $6.72 | 2 |
| THORLABS SM1T1 - SM1 (1.035"-40) Coupler, Internal Threads | [Thorlabs](https://www.thorlabs.com/thorproduct.cfm?partnumber=SM1T1) | SM1T1 | $9.80 | 1 |
| THORLABS SM1L05 - SM1 Lens Tube, 0.50" Thread Depth | [Thorlabs](https://www.thorlabs.com/thorproduct.cfm?partnumber=SM1L05E) | SM1L05 | $12.59 | 1 |
| THORLABS SM1A6 - Adapter with External SM1 Threads and Internal SM05 Threads, 0.15" Thick | [Thorlabs](https://www.thorlabs.com/thorproduct.cfm?partnumber=SM1A6) | SM1A6 | $20.17 | 1 |
| THORLABS DG10-600 - Ø1" Unmounted N-BK7 Ground Glass Diffuser, 600 Grit | [Thorlabs](https://www.thorlabs.com/thorproduct.cfm?partnumber=DG10-600) | DG10-600 | $16.13 | 1 |
| THORLABS SM1T2 SM1 (1.035"-40) Coupler, External Threads, 0.5" Long | [Thorlabs](https://www.thorlabs.com/thorproduct.cfm?partnumber=SM1T2) | SM1T2 | $20.91 | 1 |
| THORLABS SM1V10 - Ø1" Adjustable Lens Tube, 0.81" Travel Range | [Thorlabs](https://www.thorlabs.com/thorproduct.cfm?partnumber=SM1V10) | SM1V10 | $33.50 | 1 |
| THORLABS SM1A3 - Adapter with External SM1 Threads and Internal RMS Threads | [Thorlabs](https://www.thorlabs.com/thorproduct.cfm?partnumber=SM1A3) | SM1A3 | $18.50 | 1 |
| 25 mm O.D., 4 mm I.D. PLA 3D printed pinhole | Custom 3D printed. File S6a, S6b | | | 1 |
| THORLABS AC254-125-A-AC254-125-A-ML - f=125 mm, Ø1" Achromatic Doublet, SM1-Threaded Mount, ARC: 400-700 nm | [Thorlabs](https://www.thorlabs.com/thorproduct.cfm?partnumber=AC254-125-A-ML) | AC254-125-A-ML | $106.05 | 1 |
| THORLABS SM1A9 - Adapter with External C-Mount Threads and Internal SM1 Threads | [Thorlabs](https://www.thorlabs.com/thorproduct.cfm?partnumber=SM1A9) | SM1A9 | $19.38 | 1 |
| Silicone Cover Stranded-Core Wire - 2m 26AWG Black | [Adafruit](https://www.adafruit.com/product/1881) | 1881 | $0.95 | 1 |
| Silicone Cover Stranded-Core Wire - 2m 26AWG Red | [Adafruit](https://www.adafruit.com/product/2513) | 1877 | $0.95 | 1 |
| Silicone Cover Stranded-Core Wire - 2m 30AWG Black | [Adafruit](https://www.adafruit.com/product/2003) | 2003 | $0.75 | 1 |
| Silicone Cover Stranded-Core Wire - 2m 30AWG Red | [Adafruit](https://www.adafruit.com/product/2001) | 2001 | $0.75 | 1 |
| JST PH 2-Pin Cable - Female Connector 100mm | [Adafruit](https://www.adafruit.com/product/261) | 261 | $0.75 | 1 |
| JST PH 2-Pin Cable - Male Header 200mm | [Adafruit](https://www.adafruit.com/product/3814) | 3814 | $0.75 | 3 |
| 20mm Coin Cell Breakout w/On-Off Switch (CR2032) | [Adafruit](https://www.adafruit.com/product/1871) | 1871 | $3.50 | 1 |
| Rechargeable Lithium Ion 2032 Coin Cell Batteries with Battery Charger | [Amazon.com](https://www.amazon.com/CT-ENERGY-Lithium-Battery-Rechargeable-Batteries/dp/B07L933KGQ/ref=sr_1_4_sspa?crid=1CZSBFURYAI8D&dchild=1&keywords=rechargeable+2032+batteries&qid=1594036411&sprefix=rechargeable+2032+%2Caps%2C135&sr=8-4-spons&psc=1&spLa=ZW5jcnlwdGVkUXVhbGlmaWVyPUE4OVMzMUNPUERXNUUmZW5jcnlwdGVkSWQ9QTA4NjE2MzYxQVlMSlU3OFc3MEk5JmVuY3J5cHRlZEFkSWQ9QTAzNjQ3OTgySjBDWldTVVVMQzhGJndpZGdldE5hbWU9c3BfYXRmJmFjdGlvbj1jbGlja1JlZGlyZWN0JmRvTm90TG9nQ2xpY2s9dHJ1ZQ==) | NA | $23.99 | 1 |
| Cat 6 Ethernet Cable 1 Foot, Flat with Snagless RJ45 Connectors | [Amazon.com](https://www.amazon.com/Cat-Ethernet-Cable-Black-Connectors/dp/B01IQWGKQ6/ref=sr_1_1_sspa?dchild=1&keywords=1+ft+rj45+flat&qid=1594035820&sr=8-1-spons&psc=1&smid=A17K4O9J62HKE5&spLa=ZW5jcnlwdGVkUXVhbGlmaWVyPUFaQ1BCSjdHWTI4MjMmZW5jcnlwdGVkSWQ9QTAzNDEzNTgxV0MxVUZCV1hTSTROJmVuY3J5cHRlZEFkSWQ9QTA3OTU2MTRMNjI1RzJHQjUyMUcmd2lkZ2V0TmFtZT1zcF9hdGYmYWN0aW9uPWNsaWNrUmVkaXJlY3QmZG9Ob3RMb2dDbGljaz10cnVl) | NA | $12.99 | 1 |
| GPIO 8-pin 20cm JST Cable | LUCID Vision Labs | GPIO-8P20 | $10.00 | 1 |
| PLA 3D Printed Power Connector for Camera | Custom 3D printed. File S5a, S5b | | | 1 |
| PLA 3D Printed Secure GiGE Connector | This publication | This publication | NA |  |
| Lithium Ion Battery - 3.7v 2000mAh | [Adafruit](https://www.adafruit.com/product/2011) | 2011 | $12.50 | 1 |
| LED - Green with Resistor 5mm (25 pack) | [Sparkfun](https://www.sparkfun.com/products/14563) | 14563 | $8.55 | 1 |
| Multi-Colored Heat Shrink Pack - 3/32" + 1/8" + 3/16" Diameters | Adafruit | 1649 | $4.95 | 1 |
| Solder Lead Free - 100-gram Spool | [Sparkfun](https://www.sparkfun.com/products/9325) | 9325 | $7.95 | 1 |
| TP-Link AC750 Wireless Travel Router TL-WR902AC | [Amazon.com](https://www.amazon.com/TP-Link-Wireless-Travel-Router-TL-WR902AC/dp/B01N5RCZQH/ref=sr_1_2?keywords=tp-link+wr902ac&qid=1561229307&s=gateway&sr=8-2) | TL-WR902AC | $44.99 | 1 |
| Pololu 12V Step-Up Voltage Regulator U3V70F12 | [Pololu Robotics & Electronics](https://www.pololu.com/product/2895) | 2895 | $12.95 | 1 |
| Pololu 5V, Step-Up Voltage Regulator U3V70F5 | [Pololu Robotics & Electronics](https://www.pololu.com/product/2891) | 2891 | $12.95 | 1 |
| Hakko Professional Quality 20-30 AWG Wire Strippers - CSP-30-1 | [Adafruit](https://www.adafruit.com/product/527) | 527 | $14.95 | 1 |
| ATTEN 50W 110V Soldering Iron With Station - AT-937 | [Adafruit](https://www.adafruit.com/product/3565) | 3565 | $44.95 | 1 |
| Solder Vacuum | [Sparkfun](https://www.sparkfun.com/products/13203) | 13203 | $4.95 | 1 |
| THOR LABS SPW602 - Spanner Wrench for SM1-Threaded Retaining Rings, Graduated Scale with 0.1" (2.5 mm) Increments, Length = 3.88" | [Thorlabs](https://www.thorlabs.com/thorproduct.cfm?partnumber=SPW602) | SPW602 | $27.55 | 1 |
| SPW603 - Spanner Wrench for SM05-Threaded Retaining Rings, Length = 1.00" | [Thorlabs](https://www.thorlabs.com/thorproduct.cfm?partnumber=SPW603) | SPW603 | $25.24 | 1 |
| Ultimaker NFC PLA Filament 2.85mm (750g) | [3DUNIVERSE](https://shop3duniverse.com/collections/ultimaker-brand-filaments/products/ultimaker-nfc-pla-filament-2-85mm-750g) | UMNFC-PLA285-SILVER | $49.95 | 1 |
| Ultimaker NFC PVA Filament 2.85mm | [3DUNIVERSE](https://shop3duniverse.com/collections/ultimaker-brand-filaments/products/official-ultimaker-pva-filament-2-85mm) | UMPVA350 | $47.95 | 1 |
| 3D Universe LokBuild 3D Print Build Surface | [3DUNIVERSE](https://shop3duniverse.com/collections/accessories/products/3d-universe-lokbuild-3d-print-build-surface) | LOKBUILD-SM | $15.00 | 1 |
| Ultimaker 3 3D Printer | [3DUNIVERSE](https://shop3duniverse.com/collections/ultimaker-3d-printers/products/ultimaker-3) | UM3 | $3,495.00 | 1 |
| Ultimaker Cura 4.1 Software* | [Ultimaker.com](https://ultimaker.com/en/products/ultimaker-cura-software) | Ultimaker Cura 4.1.0 | $0.00 | 1 |
| Autodesk Fusion 360* | [Autodesk](https://www.autodesk.com/education/free-software/featured) | Fusion 360 Education Licence (3 yr.) | $0.00 | 1 |
| Autodesk Inventor | [Autodesk](https://www.autodesk.com/education/free-software/featured) | Inventor Professional 2020 Education License (3 yr.) | $0.00 | 1 |

**Table S2.** Resend Parameters.

| **Parameter** | **Units** | **Value** |
| --- | --- | --- |
| Resend Enable |  | True |
| Max Resend per Packet |  | 25 |
| Memory Window Size | KB | 500000 |
| Resend Threshold | % | 2 |
| Missing Packet Timeout | ms | 100 |
| New Packet Timeout | ms | 100 |
| Resend Response Timeout | ms | 10 |
| Resend Timer Resolution | ms | 10 |

**Table S3.** List of oligonucleotide sequences.

| **Name** | **Sequence** | | **Length** |
| --- | --- | --- | --- |
| **Backbone sequences (Common set of oligos for all constructs)** | | | |
| 1. 5’Biotin | (5’ 2x bio) AACATCCAATAAATCATACAGGCAAGGCAAAGAATTAGCA | | 40 |
| 2 | AAATTAAGCAATAAAGCCTC | | 20 |
| 3 | AGAGCATAAAGCTAAATCGGTTGTACCAAAAACATTATGACCCTGTAATACTTTTGCGGG | | 60 |
| 4 | AGAAGCCTTTATTTCAACGCAAGGATAAAAATTTTTAGAACCCTCATATATTTTAAATGC | | 60 |
| 5 | AATGCCTGAGTAATGTGTAGGTAAAGATTCAAAAGGGTGAGAAAGGCCGGAGACAGTCAA | | 60 |
| 6 | ATCACCATCAATATGATATTCAACCGTTCTAGCTGATAAATTAATGCCGGAGAGGGTAGC | | 60 |
| 7 | TATTTTTGAGAGATCTACAAAGGCTATCAGGTCATTGCCTGAGAGTCTGGAGCAAACAAG | | 60 |
| 8 | AGAATCGATGAACGGTAATCGTAAAACTAGCATGTCAATCATATGTACCCCGGTTGATAA | | 60 |
| 9 | TCAGAAAAGCCCCAAAAACAGGAAGATTGTATAAGCAAATATTTAAATTGTAAACGTTAA | | 60 |
| 10 | TATTTTGTTAAAATTCGCATTAAATTTTTGTTAAATCAGCTCATTTTTTAACCAATAGGA | | 60 |
| 11 | ACGCCATCAAAAATAATTCGCGTCTGGCCTTCCTGTAGCCAGCTTTCATCAACATTAAAT | | 60 |
| 12 | GTGAGCGAGTAACAACCCGTCGGATTCTCCGTGGGAACAAACGGCGGATTGACCGTAATG | | 60 |
| 13 | GGATAGGTCACGTTGGTGTAGATGGGCGCATCGTAACCGTGCATCTGCCAGTTTGAGGGG | | 60 |
| 14 | ACGACGACAGTATCGGCCTCAGGAAGATCGCACTCCAGCCAGCTTTCCGGCACCGCTTCT | | 60 |
| 15 | GGTGCCGGAAACCAGGCAAAGCGCCATTCGCCATTCAGGCTGCGCAACTGTTGGGAAGGG | | 60 |
| 16 | CGATCGGTGCGGGCCTCTTCGCTATTACGCCAGCTGGCGAAAGGGGGATGTGCTGCAAGG | | 60 |
| 17 | CGATTAAGTTGGGTAACGCCAGGGTTTTCCCAGTCACGACGTTGTAAAACGACGGCCAGT | | 60 |
| 18 | GCCAAGCTTGCATGCCTGCAGGTCGACTCTAGAGGATCCCCGGGTACCGAGCTCGAATTC | | 60 |
| 19 | GTAATCATGGTCATAGCTGTTTCCTGTGTGAAATTGTTATCCGCTCACAATTCCACACAA | | 60 |
| 20 | CATACGAGCCGGAAGCATAAAGTGTAAAGCCTGGGGTGCCTAATGAGTGAGCTAACTCAC | | 60 |
| 21 | ATTAATTGCGTTGCGCTCACTGCCCGCTTTCCAGTCGGGAAACCTGTCGTGCCAGCTGCA | | 60 |
| 22 | TTAATGAATCGGCCAACGCGCGGGGAGAGGCGGTTTGCGTATTGGGCGCCAGGGTGGTTT | | 60 |
| 23 | TTCTTTTCACCAGTGAGACGGGCAACAGCTGATTGCCCTTCACCGCCTGGCCCTGAGAGA | | 60 |
| 24 | GTTGCAGCAAGCGGTCCACGCTGGTTTGCCCCAGCAGGCGAAAATCCTGTTTGATGGTGG | | 60 |
| 25 | TTCCGAAATCGGCAAAATCCCTTATAAATCAAAAGAATAGCCCGAGATAGGGTTGAGTGT | | 60 |
| 26 | TGTTCCAGTTTGGAACAAGAGTCCACTATTAAAGAACGTGGACTCCAACGTCAAAGGGCG | | 60 |
| 27 | AAAAACCGTCTATCAGGGCGATGGCCCACTACGTGAACCATCACCCAAATCAAGTTTTTT | | 60 |
| 28 | GGGGTCGAGGTGCCGTAAAGCACTAAATCGGAACCCTAAAGGGAGCCCCCGATTTAGAGC | | 60 |
| 29 | TTGACGGGGAAAGCCGGCGAACGTGGCGAGAAAGGAAGGGAAGAAAGCGAAAGGAGCGGG | | 60 |
| 30 | CGCTAGGGCGCTGGCAAGTGTAGCGGTCACGCTGCGCGTAACCACCACACCCGCCGCGCT | | 60 |
| 31 | TAATGCGCCGCTACAGGGCGCGTACTATGGTTGCTTTGACGAGCACGTATAACGTGCTTT | | 60 |
| 32 | CCTCGTTAGAATCAGAGCGGGAGCTAAACAGGAGGCCGATTAAAGGGATTTTAGACAGGA | | 60 |
| 33 | ACGGTACGCCAGAATCCTGAGAAGTGTTTTTATAATCAGTGAGGCCACCGAGTAAAAGAG | | 60 |
| 34 | TCTGTCCATCACGCAAATTAACCGTTGTAGCAATACTTCTTTGATTAGTAATAACATCAC | | 60 |
| 35 | TTGCCTGAGTAGAAGAACTCAAACTATCGGCCTTGCTGGTAATATCCAGAACAATATTAC | | 60 |
| 36 | CGCCAGCCATTGCAACAGGAAAAACGCTCATGGAAATACCTACATTTTGACGCTCAATCG | | 60 |
| 37 | TCTGAAATGGATTATTTACATTGGCAGATTCACCAGTCACACGACCAGTAATAAAAGGGA | | 60 |
| 38 | CATTCTGGCCAACAGAGATAGAACCCTTCTGACCTGAAAGCGTAAGAATACGTGGCACAG | | 60 |
| 39 | ACAATATTTTTGAATGGCTATTAGTCTTTAATGCGCGAACTGATAGCCCTAAAACATCGC | | 60 |
| 40 | CATTAAAAATACCGAACGAACCACCAGCAGAAGATAAAACAGAGGTGAGGCGGTCAGTAT | | 60 |
| 41 | TAACACCGCCTGCAACAGTGCCACGCTGAGAGCCAGCAGCAAATGAAAAATCTAAAGCAT | | 60 |
| 42 | CACCTTGCTGAACCTCAAATATCAAACCCTCAATCAATATCTGGTCAGTTGGCAAATCAA | | 60 |
| 43 | CAGTTGAAAGGAATTGAGGAAGGTTATCTAAAATATCTTTAGGAGCACTAACAACTAATA | | 60 |
| 44 | GATTAGAGCCGTCAATAGATAATACATTTGAGGATTTAGAAGTATTAGACTTTACAAACA | | 60 |
| 45 | ATTCGACAACTCGTATTAAATCCTTTGCCCGAACGTTATTAATTTTAAAAGTTTGAGTAA | | 60 |
| 46 | CATTATCATTTTGCGGAACAAAGAAACCACCAGAAGGAGCGGAATTATCATCATATTCCT | | 60 |
| 47 | GATTATCAGATGATGGCAATTCATCAATATAATCCTGATTGTTTGGATTATACTTCTGAA | | 60 |
| 48 | TAATGGAAGGGTTAGAACCTACCATATCAAAATTATTTGCACGTAAAACAGAAATAAAGA | | 60 |
| 49 | AATTGCGTAGATTTTCAGGTTTAACGTCAGATGAATATACAGTAACAGTACCTTTTACAT | | 60 |
| 50 | CGGGAGAAACAATAACGGATTCGCCTGATTGCTTTGAATACCAAGTTACAAAATCGCGCA | | 60 |
| 51 | GAGGCGAATTATTCATTTCAATTACCTGAGCAAAAGAAGATGATGAAACAAACATCAAGA | | 60 |
| 52 | AAACAAAATTAATTACATTTAACAATTTCATTTGAATTACCTTTTTTAATGGAAACAGTA | | 60 |
| 53 | CATAAATCAATATATGTGAGTGAATAACCTTGCTTCTGTAAATCGTCGCTATTAATTAAT | | 60 |
| 54 | TTTCCCTTAGAATCCTTGAAAACATAGCGATAGCTTAGATTAAGACGCTGAGAAGAGTCA | | 60 |
| 55 | ATAGTGAATTTATCAAAATCATAGGTCTGAGAGACTACCTTTTTAACCTCCGGCTTAGGT | | 60 |
| 56 | TGGGTTATATAACTATATGTAAATGCTGATGCAAATCCAATCGCAAGACAAAGAACGCGA | | 60 |
| 57 | GAAAACTTTTTCAAATATATTTTAGTTAATTTCATCTTCTGACCTAAATTTAATGGTTTG | | 60 |
| 58 | AAATACCGACCGTGTGATAAATAAGGCGTTAAATAAGAATAAACACCGGAATCATAATTA | | 60 |
| 59 | CTAGAAAAAGCCTGTTTAGTATCATATGCGTTATACAAATTCTTACCAGTATAAAGCCAA | | 60 |
| 60 | CGCTCAACAGTAGGGCTTAATTGAGAATCGCCATATTTAACAACGCCAACATGTAATTTA | | 60 |
| 61 | GGCAGAGGCATTTTCGAGCCAGTAATAAGAGAATATAAAGTACCGACAAAAGGTAAAGTA | | 60 |
| 62 | ATTCTGTCCAGACGACGACAATAAACAACATGTTCAGCTAATGCAGAACGCGCCTGTTTA | | 60 |
| 63 | TCAACAATAGATAAGTCCTGAACAAGAAAAATAATATCCCATCCTAATTTACGAGCATGT | | 60 |
| 64 | AGAAACCAATCAATAATCGGCTGTCTTTCCTTATCATTCCAAGAACGGGTATTAAACCAA | | 60 |
| 65 | GTACCGCACTCATCGAGAACAAGCAAGCCGTTTTTATTTTCATCGTAGGAATCATTACCG | | 60 |
| 66 | CGCCCAATAGCAAGCAAATCAGATATAGAAGGCTTATCCGGTATTCTAAGAACGCGAGGC | | 60 |
| 67 | GTTTTAGCGAACCTCCCGACTTGCGGGAGGTTTTGAAGCCTTAAATCAAGATTAGTTGCT | | 60 |
| 68 | ATTTTGCACCCAGCTACAATTTTATCCTGAATCTTACCAACGCTAACGAGCGTCTTTCCA | | 60 |
| 69 | GAGCCTAATTTGCCAGTTACAAAATAAACAGCCATATTATTTATCCCAATCCAAATAAGA | | 60 |
| 70 | AACGATTTTTTGTTTAACGTCAAAAATGAAAATAGCAGCCTTTACAGAGAGAATAACATA | | 60 |
| 71 | AAAACAGGGAAGCGCATTAGACGGGAGAATTAACTGAACACCCTGAACAAAGTCAGAGGG | | 60 |
| 72 | TAATTGAGCGCTAATATCAGAGAGATAACCCACAAGAATTGAGTTAAGCCCAATAATAAG | | 60 |
| 73 | AGCAAGAAACAATGAAATAGCAATAGCTATCTTACCGAAGCCCTTTTTAAGAAAAGTAAG | | 60 |
| 74 | CAGATAGCCGAACAAAGTTACCAGAAGGAAACCGAGGAAACGCAATAATAACGGAATACC | | 60 |
| 75 | CAAAAGAACTGGCATGATTAAGACTCCTTATTACGCAGTATGTTAGCAAACGTAGAAAAT | | 60 |
| 76 | ACATACATAAAGGTGGCAACATATAAAAGAAACGCAAAGACACCACGGAATAAGTTTATT | | 60 |
| 77 | TTGTCACAATCAATAGAAAATTCATATGGTTTACCAGCGCCAAAGACAAAAGGGCGACAT | | 60 |
| 78 | TCAACCGATTGAGGGAGGGAAGGTAAATATTGACGGAAATTATTCATTAAAGGTGAATTA | | 60 |
| 79 | TCACCGTCACCGACTTGAGCCATTTGGGAATTAGAGCCAGCAAAATCACCAGTAGCACCA | | 60 |
| 80 | TTACCATTAGCAAGGCCGGAAACGTCACCAATGAAACCATCGATAGCAGCACCGTAATCA | | 60 |
| 81 | GTAGCGACAGAATCAAGTTTGCCTTTAGCGTCAGACTGTAGCGCGTTTTCATCGGCATTT | | 60 |
| 82 | TCGGTCATAGCCCCCTTATTAGCGTTTGCCATCTTTTCATAATCAAAATCACCGGAACCA | | 60 |
| 83 | GAGCCACCACCGGAACCGCCTCCCTCAGAGCCGCCACCCTCAGAACCGCCACCCTCAGAG | | 60 |
| 84 | CCACCACCCTCAGAGCCGCCACCAGAACCACCACCAGAGCCGCCGCCAGCATTGACAGGA | | 60 |
| 85 | GGTTGAGGCAGGTCAGACGATTGGCCTTGATATTCACAAACAAATAAATCCTCATTAAAG | | 60 |
| 86 | CCAGAATGGAAAGCGCAGTCTCTGAATTTACCGTTCCAGTAAGCGTCATACATGGCTTTT | | 60 |
| 87 | GATGATACAGGAGTGTACTGGTAATAAGTTTTAACGGGGTCAGTGCCTTGAGTAACAGTG | | 60 |
| 88 | CCCGTATAAACAGTTAATGCCCCCTGCCTATTTCGGAACCTATTATTCTGAAACATGAAA | | 60 |
| 89 | GTATTAAGAGGCTGAGACTCCTCAAGAGAAGGATTAGGATTAGCGGGGTTTTGCTCAGTA | | 60 |
| 90 | CCAGGCGGATAAGTGCCGTCGAGAGGGTTGATATAAGTATAGCCCGGAATAGGTGTATCA | | 60 |
| 91 | CCGTACTCAGGAGGTTTAGTACCGCCACCCTCAGAACCGCCACCCTCAGAACCGCCACCC | | 60 |
| 92 | TCAGAGCCACCACCCTCATTTTCAGGGATAGCAAGCCCAATAGGAACCCATGTACCGTAA | | 60 |
| 93 | CACTGAGTTTCGTCACCAGTACAAACTACAACGCCTGTAGCATTCCACAGACAGCCCTCA | | 60 |
| 94 | TAGTTAGCGTAACGATCTAAAGTTTTGTCGTCTTTCCAGACGTTAGTAAATGAATTTTCT | | 60 |
| 95 | GTATGGGATTTTGCTAAACAACTTTCAACAGTTTCAGCGGAGTGAGAATAGAAAGGAACA | | 60 |
| 96 | ACTAAAGGAATTGCGAATAATAATTTTTTCACGTTGAAAATCTCCAAAAAAAAGGCTCCA | | 60 |
| 97 | AAAGGAGCCTTTAATTGTATCGGTTTATCAGCTTGCTTTCGAGGTGAATTTCTTAAACAG | | 60 |
| 98 | CTTGATACCGATAGTTGCGCCGACAATGACAACAACCATCGCCCACGCATAACCGATATA | | 60 |
| 99 | TTCGGTCGCTGAGGCTTGCAGGGAGTTAAAGGCCGCTTTTGCGGGATCGTCACCCTCAGC | | 60 |
| 100 | AGCGAAAGACAGCATCGGAACGAGGGTAGCAACGGCTACAGAGGCTTTGAGGACTAAAGA | | 60 |
| 101 | CTTTTTCATGAGGAAGTTTCCATTAAACGGGTAAAATACGTAATGCCACTACGAAGGCAC | | 60 |
| 102 | CAACCTAAAACGAAAGAGGCAAAAGAATACACTAAAACACTCATCTTTGACCCCCAGCGA | | 60 |
| 103 | TTATACCAAGCGCGAAACAAAGTACAACGGAGATTTGTATCATCGCCTGATAAATTGTGT | | 60 |
| 104 | CGAAATCCGCGACCTGCTCCATGTTACTTAGCCGGAACGAGGCGCAGACGGTCAATCATA | | 60 |
| 105 | AGGGAACCGAACTGACCAACTTTGAAAGAGGACAGATGAACGGTGTACAGACCAGGCGCA | | 60 |
| 106 | TAGGCTGGCTGACCTTCATCAAGAGTAATCTTGACAAGAACCGGATATTCATTACCCAAA | | 60 |
| 107 | TCAACGTAACAAAGCTGCTCATTCAGTGAATAAGGCTTGCCCTGACGAGAAACACCAGAA | | 60 |
| 108 | CGAGTAGTAAATTGGGCTTGAGATGGTTTAATTTCAACTTTAATCATTGTGAATTACCTT | | 60 |
| 109 | ATGCGATTTTAAGAACTGGCTCATTATACCAGTCAGGACGTTGGGAAGAAAAATCTACGT | | 60 |
| 110 | TAATAAAACGAACTAACGGAACAACATTATTACAGGTAGAAAGATTCATCAGTTGAGATT | | 60 |
| 111 | TAGGAATACCACATTCAACTAATGCAGATACATAACGCCAAAAGGAATTACGAGGCATAG | | 60 |
| 112 | TAAGAGCAACACTATCATAACCCTCGTTTACCAGACGACGATAAAAACCAAAATAGCGAG | | 60 |
| 113 | AGGCTTTTGCAAAAGAAGTTTTGCCAGAGGGGGTAATAGTAAAATGTTTAGACTGGATAG | | 60 |
| 114 | CGTCCAATACTGCGGAATCGTCATAAATATTCATTGAATCCCCCTCAAATGCTTTAAACA | | 60 |
| 115 | GTTCAGAAAACGAGAATGACCATAAATCAAAAATCAGGTCTTTACCCTGACTATTATAGT | | 60 |
| 116 | CAGAAGCAAAGCGGATTGCATCAAAAAGATTAAGAGGAAGCCCGAAAGACTTCAAATATC | | 60 |
| 117 | GCGTTTTAATTCGAGCTTCAAAGCGAACCAGACCGGAAGCAAACTCCAACAGGTCAGGAT | | 60 |
| 118 | TAGAGAGTACCTTTAATTGCTCCTTTTGATAAGAGGTCATTTTTGCGGATGGCTTAGAGC | | 60 |
| 119 | TTAATTGCTGAATATAATGCTGTAGCTCAACATGTTTTAAATATGCAACTAAAGTACGGT | | 60 |
| 120 | GTCTGGAAGTTTCATTCCATATAACAGTTGATTCCCAATTCTGCGAACGAGTAGATTTAG | | 60 |
| 121 | TTTGACCATTAGATACATTTCGCAAATGGTCAATAACCTGTTTAGCTAT | | 49 |
| 122 | ATTTTCATTTGGGGCGCGAGCTGAAAAGGT | | 30 |
| **Variable Overhanging sequences specific for each construct (overhanging regions underlined)** | | | |
| 123.1 - 7nt GC 71 - Over | | GGCATCAATTCTACTAATAGTAGTAGCATTGCAACGG |  |
| 123.1 - 7nt GC 71 - Comp | | GGCATCAATTCTACTAATAGTAGTAGCATTCCGTTGC |  |
| 123.1 - 7nt GC 57 - Over | | GGCATCAATTCTACTAATAGTAGTAGCATTGCAAAGG |  |
| 123.1 - 7nt GC 57 - Comp | | GGCATCAATTCTACTAATAGTAGTAGCATTCCTTTGC |  |
| 123.1 - 8nt - Over | | GGCATCAATTCTACTAATAGTAGTAGCATTGCAAGCGG |  |
| 123.1 - 8nt - Comp | | GGCATCAATTCTACTAATAGTAGTAGCATTCCGCTTGC |  |
| 123.1 - 9nt - Over | | GGCATCAATTCTACTAATAGTAGTAGCATTGTCAAGCGG |  |
| 123.1 - 9nt - Comp | | GGCATCAATTCTACTAATAGTAGTAGCATTCCGCTTGAC |  |

**Table S4:** Mass calculation for micro-spheres.

| **Bead** | **M270** | **SVP-50-5** |
| --- | --- | --- |
| **Radius (µm)** | 1.40 ± 0.09 | 2.51 ± 0.25 |
| **Density (g/cm)** | 1.61± 0.02 | 1.05 ± 0.01 |
| **Density of 1X PBS (g/cm)** | 1.01± 0.01 | 1.01 ± 0.01 |
| **Effective density (g/cm)** | 0.60 ± 0.02 | 0.04 ± 0.01 |
| **Effective mass of bead in 1X PBS (g)** | 6.9E-12 ± 8.0*10^-13 | 2.6*10^-12 ± 7.6*10^-13 |

**Table S5:** RPM used to achieve the prescribed force for M270 and SVP-50-5 beads.

| **Force** | **M270 (RPM)** | **SVP-50-5 (RPM)** |
| --- | --- | --- |
| 12 | 1093 |  |
| 10 | 997 |  |
| 8 | 892 |  |
| 6 | 773 |  |
| 4.5 |  | 1080 |
| 4 | 630 |  |
| 2 | 446 |  |
