## Supplementary material for "A Wi-Fi live streaming Centrifuge Force Microscope for benchtop single-molecule experiments": 8. Bead dissociation analysis.m

```
clear all
close all
%% %% load in video and accessory files
%%%%%%%%%%%%%%%%%%%%%%%%%%%%%%%%%%%%%%%%%%%%%%%%%%%%%%%%%%%%%%%%%%%%%%%%%
%Select folder containing images, select starting and ending frames, read
%timestamp file
%%%%%%%%%%%%%%%%%%%%%%%%%%%%%%%%%%%%%%%%%%%%%%%%%%%%%%%%%%%%%%%%%%%%%%%%%
[filename,pathname] = uigetfile('*.*','Select first movie frame');
cd (pathname);
separators = strfind(pathname, '\');
movie_name = pathname(separators(end-1)+1:separators(end)-1);
files = dir('*.tiff');

%re-sort the files into correct numerical order
filenames = {files.name};
str  = sprintf('%s#', filenames{:});
num  = sscanf(str, '%d.tiff#');
[dummy, index] = sort(num);
filenames = filenames(index);

%read timestamp file
fileID = fopen('timestamp.txt');
timestamp = fscanf(fileID, '%f ', [1 inf]);
timestamp=timestamp';

%Prompt for entering starting and ending frames
total_frames = size(files,1);
prompt = 'Start Frame:';
start_frame = input(prompt);
prompt = 'End_Frame:';
end_frame = input(prompt);

%%
%%%%%%%%%%%%%%%%%%%%%%%%%%%%%%%%%%%%%%%%%%%%%%%%%%%%%%%%%%%%%
%%Find number of initial beads & Identifing the useable beads
%%%%%%%%%%%%%%%%%%%%%%%%%%%%%%%%%%%%%%%%%%%%%%%%%%%%%%%%%%%%%

%Read the first frame after centrifuge assumes it intended speed
First_image=imread(filenames{start_frame});
video_height = size(First_image,1);
video_width = size(First_image,2);

%If the first frame has missing packets, it will mess with circle finding
%Algorithm. This section will help to crop the first image without missing
%packets which can be used as an input parameter to circle finding algoritm

%if ( (all(First_image(:))>0) == 0)     
figure(1)
imshow(First_image, 'InitialMagnification', 30)
title('Choose the region of Interest','FontSize',18, 'FontName','Arial', 'FontWeight','bold')
ROItempImg = imcrop;  
%end

%Bead Identification sensitivity hint and finetuning
SenIndex = [0.8, 0.85, 0.9];
for counter1=1:3
    [centers,radius] = imfindcircles(ROItempImg,[10 30], 'ObjectPolarity','dark','Sensitivity', SenIndex(counter1));
    figname = "Bead Identification Sensitivity-%d";
    figname=sprintf(figname,SenIndex(counter1));
    figure ('name', figname, 'NumberTitle', 'off');
    imshow(First_image, 'InitialMagnification', 45);
    viscircles(centers, radius,'EdgeColor','b','LineWidth',0.5);
    clear centers;
    clear radius;
end
prompt = 'If you want to finetune sensitivity further, enter Y/N:';
finetune = input(prompt,'s');
while (finetune == 'Y' || finetune == 'y') 
      clear SenIndex;
      prompt = 'Enter Sensitivity Index (0-1):';
      SenIndex = input(prompt);
      while (SenIndex > 1 || SenIndex <0 )
          prompt = 'Not a valid Input. Enter a value between 0 and 1:';
          SenIndex = input(prompt);
      end
      [centers,radius] = imfindcircles(ROItempImg,[10 30], 'ObjectPolarity','dark','Sensitivity', SenIndex);
      figname = "Bead Identification Sensitivity-%d";
      figname=sprintf(figname,SenIndex);
      figure ('name', figname, 'NumberTitle', 'off');
      imshow(First_image, 'InitialMagnification', 50);
      viscircles(centers, radius,'EdgeColor','b','LineWidth',0.5);
      clear finetune;
      prompt = 'Do you  want to finetune sensitivity further, enter F , else enter E:';
      finetune = input(prompt,'s');
      close;
end

clear counter1;
clear SenIndex;
clear centers;
clear radius;
close all;
prompt = 'Enter Your Preffered Sensitivity Index (0-1):';
SenIndex = input(prompt);
[centers,radius] = imfindcircles(ROItempImg,[10 30], 'ObjectPolarity','dark','Sensitivity', SenIndex);
num_total_beads = length(centers);

% Good Refrence Bead Selector
figure(1)
imshow(First_image, 'InitialMagnification', 50)
title('Choose a single tetherd bead for your reference','FontSize',18, 'FontName','Arial', 'FontWeight','bold')
% Choosing reference bead to compare with individual bead
J = imcrop;
figure(2)
imshow(J,'InitialMagnification', 250) 
[cent_ref_1,rad_ref_1] = imfindcircles(J,[10 30], 'ObjectPolarity','dark','Sensitivity', SenIndex);
roi_square_radius = 35;
roi_width = 2*roi_square_radius+1;
left1 = round(cent_ref_1(1,1))-roi_square_radius;
bottom1 = round(cent_ref_1(1,2))-roi_square_radius;
J = imcrop(J, [left1 bottom1 roi_width  roi_width]) ;

% Automatic Unuseable Bead Identifier
% Identifier for beads on the edge of the frame
for counter1=1:num_total_beads
    
    roi_square_radius = 31;
    roi_width = 2*roi_square_radius+1;
    left1 = round(centers(counter1,1))-roi_square_radius;
    bottom1 = round(centers(counter1,2))-roi_square_radius;
    tempimg= imcrop(First_image(:,:,1), [left1 bottom1 roi_width  roi_width]) ;
    [sq1,sq2]=size(tempimg);

    if sq1==sq2 && sq1>61
         beadquality(counter1)="Single";
    else
        beadquality(counter1)="Multiple";
    end
    clear tempimg; 
end 
clear counter1;

%Identifier for very close beads
for counter1=1:num_total_beads   
   for counter2=1:num_total_beads
       if counter1~=counter2
            d=sqrt((centers(counter1,1)-centers(counter2,1))^2 + (centers(counter1,2)-centers(counter2,2))^2);
           if d<40           %previously 35
            beadquality(counter1)="Multiple";
            beadquality(counter2)="Multiple";
           end
       end    
   end
end

clear counter1;
clear counter2;

% Sorting rest of the beads to Single tethered (Useable) or unuseable from
%the user end
figure(3)
imshow(First_image, 'InitialMagnification', 55);
title('Reference Frame','FontSize',18, 'FontName','Arial', 'FontWeight','bold');
for counter1=1:num_total_beads
   x = centers(counter1,1);
   y = centers(counter1,2);
   r = radius(counter1)+10;
   ang=0:0.01:2*pi; 
   xp=r*cos(ang);
   yp=r*sin(ang);
   
   hold on 
     if beadquality(counter1)=="Multiple"
       plot(x+xp,y+yp, 'r')    
   end       
   end

hold off

figure(3)
movegui(3,'northeast');
hold on
counter1=1;
counter2=1;
while counter1<=num_total_beads
    if beadquality(counter1)=="Single"
    roi_square_radius = 31;
    roi_width = 2*roi_square_radius+1;
    left1 = round(centers(counter1,1))-roi_square_radius;
    bottom1 = round(centers(counter1,2))-roi_square_radius;
    K = imcrop(First_image, [left1 bottom1 roi_width  roi_width]) ;
    sizeJ=size(J);
    sizeK=size(K);
    h=min( sizeJ(1,1), sizeK(1,1));
    w=min(sizeJ(1,2), sizeK(1,2));
    J1=J(1:h,1:w);
    K1=K(1:h,1:w);
    newImg = [J1, K1];
    figure (2)
    movegui(2,'northwest');
    imshow (newImg,'InitialMagnification', 300)
    title('Reference Bead                    Current Bead ','FontSize',10, 'FontName','Arial', 'FontWeight','bold')


       c1(1,1)=round(sizeJ(1,1)/2);
       c1_1 = J(c1(1,1),:);
       c1_1=c1_1-min(c1_1);
       c2(1,1)=round(sizeK(1,1)/2);
       c2_1 = K(c2(1,1),:);  
       c2_1=c2_1-min(c2_1);
              
       figure (4)
       movegui(4,'southwest');
       ay1 = subplot(3,1,1);
       plot (ay1,c1_1);
       title(ay1,'Reference Bead Profile');
       ay2 = subplot(3,1,2);
       plot (ay2,c2_1, 'r');
       title(ay2,'Current Bead Profile');
       ay3 = subplot(3,1,3);
       plot (ay3,c1_1);
       hold on
       plot (ay3,c2_1, 'r');
       title(ay3,'Current Bead Profile');
       hold off
 
   x = centers(counter1,1);
   y = centers(counter1,2);
   r = radius(counter1)+10;
   ang=0:0.01:2*pi; 
   xp=r*cos(ang);
   yp=r*sin(ang);
   figure(3)
   plot(x+xp,y+yp, 'y --');
   prompt = 'Single Y/N:';
   single = input(prompt,'s');
   close 2;
   close 4;
    while (single ~= "Y" & single ~= "y" & single ~= "N" & single ~= "n")
          figure(3)
          prompt = 'Not a valid Input. Single Y/N:';
          single = input(prompt,'s');
    end
   if single== "Y" | single== "y" 
       figure(3)
       plot(x+xp,y+yp, 'g')
       centersingle(counter2,:)=centers(counter1,:);
       radiusingle(counter2)= radius(counter1);
       counter2=counter2+1;
       beadquality(counter1)="Single";
       
       elseif single== "N" | single== "n" 
              figure(3)
              plot(x+xp,y+yp, 'r')
              beadquality(counter1)="Multiple";
    end
  
    clear c1;
    clear c2;
    clear c1_1;
    clear c2_1;
    clear r1;
    clear r2;
    end  
     counter1=counter1+1;
end

hold off
figure (3);
saveas(3, 'Refrence Frame', 'tif');
close 3;
close 1;
single_tether_beads = length(centersingle); 
clear counter1;
clear counter2;
clear ROItempImg;
clear finetune;


%% 
%%%%%%%%%%%%%%%%%%%%%%%%%%%%%%%%%%%%%%%%%%%%%%%%%%%%%%%%%%%%%%%%%%%%%%%%%%%%%%%%%%%%%%%%%%%%%%%%%%%%%%%%%%%%%%%%%%%%%%%%%
%%cropping bead images into a 4D matrix where 2D images of each bead(3rd dimention) is stored in the 4th dimention (time)
%%%%%%%%%%%%%%%%%%%%%%%%%%%%%%%%%%%%%%%%%%%%%%%%%%%%%%%%%%%%%%%%%%%%%%%%%%%%%%%%%%%%%%%%%%%%%%%%%%%%%%%%%%%%%%%%%%%%%%%%%

%Time resolution of data collection and analysis
ExpTimeRes = mean(diff(timestamp))       %% The average time resolution of the data
prompt = 'Enter the Time Resolution for your analysis in multiples of ExpTimeRes (in seconds):';
time_res = input(prompt);
time_res=round(time_res/ExpTimeRes);

%%Images are read into the memory prior to cropping to ROI 
totalframe = end_frame-start_frame;
for counter3=1:(totalframe/time_res) 
    if (counter3==1)
         filetoread = start_frame;  
    else    
         filetoread=start_frame+ (counter3*time_res);
    end
     chamberimage(:,:,counter3)=imread(filenames{filetoread});
end
clear counter3;

% Cropping The ROI and saving in time-matrix
counter3=1;
for counter4=1:num_total_beads
     if beadquality(counter4)=="Single"
       roi_square_radius = 31;
       roi_width = 2*roi_square_radius+1;
       left1 = round(centers(counter4,1))-roi_square_radius;
       bottom1 = round(centers(counter4,2))-roi_square_radius;
          for counter5=1:(totalframe/time_res)
            tempimg= imcrop(chamberimage(:,:,counter5), [left1 bottom1 roi_width  roi_width]) ;
            beadIM(:,:,counter3,counter5)=tempimg;
            beadIM_TC(:,:,counter3,counter5)=tempimg;
% Correction for transmission loss correction        
            if ( (all(tempimg(:))>0) == 0 && counter5>1)
                 beadIM_TC(:,:,counter3,counter5)= beadIM_TC(:,:,counter3,counter5-1);
            end
     
            clear tempimg;
          end
        counter3=counter3+1;
    end
end
clear counter3;
clear counter4;
clear counter5;
clear chamberimage;

%% Creation of bead kymograp 

for counter6=1:single_tether_beads
     
        beadkymo=beadIM(:,:,counter6,1);
        beadkymo_TC=beadIM_TC(:,:,counter6,1);  %for Transmission loss Correction
        for counter7=2:(totalframe/time_res)
            beadkymo=horzcat(beadkymo,beadIM(:,:,counter6,counter7));
            beadkymo_TC=horzcat(beadkymo_TC,beadIM_TC(:,:,counter6,counter7));   %for Transmission loss Correction
        end
        beadkymoarray(:,:,counter6)=beadkymo;
        beadkymoarray_TC(:,:,counter6)=beadkymo_TC;
        
        clear beadkymo;
   
end 

%saving bead profile across time for individual beads - The transmission
%corrected images are saved

for counter8=1:single_tether_beads
        framename = "Single tethered Bead # %d";
        framename=sprintf(framename,counter8);
        beadkymofig=figure ('Name', framename,'NumberTitle','off');
        imshow (beadkymoarray_TC(:,:,counter8));
        saveas(beadkymofig, framename, 'tif');
        close; 
        clear beadkymofig;
end
%% Stacking the bead kymographs
counter10=1;
counter11=1;
    DataIDtop=[1:20:10000];
    DataIDbottom=[20:20:10000];
for counter9=1:single_tether_beads
        if counter10==1
           TempImg= beadkymoarray_TC(:,:,counter9); 
          else
           TempImg = vertcat(TempImg, beadkymoarray_TC(:,:,counter9));  
        end   
   
    if mod(counter9,20)==0
       All_Single_Beads_TC(:,:,counter11)=TempImg;
       figname = "Beads from %d to %d";
       figname=sprintf(figname,DataIDtop(counter11),DataIDbottom(counter11));
       stackkymo =  figure ('name', figname, 'NumberTitle', 'off');
       imshow (All_Single_Beads_TC(:,:,counter11)); 
       saveas(stackkymo,figname,'tif');
       counter11=counter11+1;
       counter10=1;
    elseif (counter9==single_tether_beads)
       All_Single_Beads_TC_Last_Few(:,:,1)=TempImg; 
       DataIDbottom(counter11)=single_tether_beads;
       figname = "Beads from %d to %d";
       figname=sprintf(figname,DataIDtop(counter11),DataIDbottom(counter11));
       stackkymo =  figure ('name', figname, 'NumberTitle', 'off');
       imshow (All_Single_Beads_TC_Last_Few(:,:,1)); 
       saveas(stackkymo,figname,'tif');
    else
        counter10=0;
    end     

end
 
clear counter6;
clear counter7;
clear counter8;
clear counter9;
clear counter10;

%% Variance profile for all  beads
for counter11=1:single_tether_beads
        for counter12 = 1:(totalframe/time_res)
            tempImstore = beadIM_TC(:,:,counter11,counter12);
            tempImstore = reshape (tempImstore(:,:),[],1);
            tempImstore=im2double(tempImstore);
            varprofile(counter11,counter12)= var(tempImstore);
            clear tempImstore;
        end  
end

%saving the bead profile for indivivual beads

clear counter11;
clear counter12;
clear textname;
clear textname1;
clear textname2;
clear textname3;

%% dissociation time analysis
        
for counter14=1:single_tether_beads
        diffprof = diff(varprofile(counter14,:));
        diffprof=abs(diffprof);
        counter15=0;                                     %% Conditional counter for transferring the information that a bead dissociated or not
        for counter16=1:(totalframe/time_res)-2
            %if counter16==1
               varianceindex=10* mean(diffprof);
             % else   
              % varianceindex=30*(diffprof(counter16-1)+diffprof(counter16+1));
            %end   
            if (diffprof(counter16) > varianceindex)     %%diffprof(counter16)>(mean(diffprof)- 5*std(diffprof))     %%finetune the sensitivity here for bead dissociation algorithm
               counter15=counter15+1; 
               dissframe(counter14)= counter16;
               disstime=timestamp(start_frame+((counter16)*time_res))-timestamp(start_frame);
               timefolder(counter15)=disstime;
               peakfolder(counter15)=diffprof(counter16);
            end
        end
        if counter15==1                                  %%means bead dissociated
           beaddisstime(counter14)= disstime;
           elseif counter15<1
                  beaddisstime(counter14)=  100000;       %% means beads never disociated
        else
             [peakmax,timeIndex]=max(peakfolder);                                            %%beaddisstime(counter14)=  100001;  %% means more than one dissocation even accounted foe which is impossible.Manually correct
             disstime=timefolder(timeIndex);
             beaddisstime(counter14)= disstime;
        end
   clear timefolder;
   clear peakfolder;
   clear peakmax;
   clear timeIndex;
end   
beaddisstime=beaddisstime';
clear counter14;
clear counter15;
clear counter16;

%%
beaddisstime=beaddisstime';
%%
```
